# Nuclear Pyruvate Dehydrogenase Complex Regulates Histone Acetylation and Transcriptional Regulation in the Ethylene Response

**DOI:** 10.1101/2023.10.25.564010

**Authors:** Zhengyao Shao, Liangqiao Bian, Shyon K. Ahmadi, Tyler J. Daniel, Miguel A. Belmonte, Jackson G. Burns, Prashanth Kotla, Yang Bi, Zhouxin Shen, Shou-Ling Xu, Zhi-Yong Wang, Steven P. Briggs, Hong Qiao

## Abstract

Ethylene plays its essential roles in plant development, growth, and defense responses by controlling the transcriptional reprograming, in which EIN2-C-directed regulation of histone acetylation is the first key-step for chromatin to perceive ethylene signaling. But how the nuclear acetyl coenzyme A (acetyl CoA) is produced to ensure the ethylene-mediated histone acetylation is unknown. Here we report that ethylene triggers the accumulation of the pyruvate dehydrogenase complex (PDC) in the nucleus to synthesize nuclear acetyl CoA to regulate ethylene response. PDC is identified as an EIN2-C nuclear partner, and ethylene triggers its nuclear accumulation. Mutations in PDC lead to an ethylene-hyposensitivity that results from the reduction of histone acetylation and transcription activation. Enzymatically active nuclear PDC synthesize nuclear acetyl CoA for EIN2-C-directed histone acetylation and transcription regulation. These findings uncover a mechanism by which PDC-EIN2 converges the mitochondrial enzyme mediated nuclear acetyl CoA synthesis with epigenetic and transcriptional regulation for plant hormone response.

## Introduction

Ethylene is a pivotal plant hormone that plays pleiotropic roles in plant growth, development, stress response and defense response to pathogens by controlling downstream gene transcription, protein translation and posttranslational modifications, mitochondrial retrograde signaling, chromatin remodeling, and epigenetic regulation of gene expression^1–14^. Ethylene signal is perceived by ethylene receptors embedded on the endoplasmic reticulum (ER) membrane^15,16^. In the absence of ethylene, both the ethylene receptors and CONSTITUTIVE TRIPLE RESPONSE 1 (CTR1), an ER membrane associated kinase that directly interacts with one of ethylene receptors ETHYLENE RESPONSE1 (ETR1), are activated^17,18^. CTR1 phosphorylates ETHYLENE INSENSTIVE 2 (EIN2), the essential positive regulator of ethylene signaling, at its C-terminal end (EIN2-C), leading to the repression of the EIN2 activity^19,20^. Without ethylene signaling activation, EIN2 is localized to the ER membrane at which it interacts with two F-box proteins EIN2 TARGETING PROTEIN 1/2 (ETP1/2) that mediate its protein degradation via the ubiquitin-proteasome pathway^21^. Upon the perception of ethylene, both ETR1 and CTR1 are inactivated; the EIN2-C is dephosphorylated through an unknown mechanism^20^. The dephosphorylated EIN2-C is cleaved and translocated into both the nucleus and the P-body^20,22,23^. In the P-body, EIN2-C mediates the translational repression of two F-box proteins EIN3-BINDING F BOX PROTEIN 1/2 (EBF1/2) to promote the protein accumulation of ETHYLENE INSENSITIVE 3 (EIN3), the key transcription activator that is sufficient and necessary for activation of all ethylene-response genes^6,23,24^. In the nucleus, EIN2-C mediates the direct regulation of histone acetylation of H3K14 and H3K23 via histone binding protein EIN2 NUCLEAR ASSOCIATED PROTEIN 1/2 (ENAP1/2), leading to an EIN3 dependent transcriptional regulation for ethylene response^25–27^.

Epigenetic regulation of gene expression plays critical roles in various plant developmental processes and stress response^28–32^. Histone acetylation promotes the relaxation of nucleosome between wrapped DNA and histone octamer by neutralizing the positive charges of lysine residues in histones, which is crucial for all DNA-based processes, including DNA replication and gene transcription^33,34^. Histone acetyltransferases (HATs) transfer acetyl group from acetyl coenzyme A (acetyl CoA) to conserved lysine residues on histone N-terminal tails to acetylate histones. In eukaryotes, the biosynthesis of acetyl CoA is thought to occur in the subcellular compartment where it is required because of its membrane impermeability and high instability due to the high-energy thioester bond that joins the acetyl and CoA groups^35^. The availability and abundance of acetyl CoA are therefore crucial for HAT enzymatic activity and histone acetylation. For most HATs, their Michaelis constants (Kms), the concentration of substrate which permits the enzyme to achieve half Vmax, lie in the range within or greater than the cellular acetyl CoA concentrations in order to sense and respond to the fluctuation of acetyl CoA production^36^.

Therefore, the levels of histone acetylation are often linked with the availability of acetyl CoA. Such metabolic regulation of histone acetylation by nuclear synthesis of acetyl CoA has been reported to govern many major biological events including cell proliferation, cell fate determination, and environmental response^37–43^. However, the relationship between the nuclear acetyl CoA and histone acetylation in plant hormone response is completely unknown.

In this study, we report that functional pyruvate dehydrogenase complex (PDC) can translocate from the mitochondria to the nucleus in the presence of ethylene, generating a nuclear pool of acetyl CoA from pyruvate for the EIN2-directed acetylation of core histones for transcriptional regulation in the ethylene response. In the presence of ethylene, the PDC complex translocates to the nucleus where it interacts with EIN2-C. Mutations in PDC lead to a decreased ethylene response at both at genetic level and transcriptional level. ChIP-seq assay reveals that ethylene-induced elevation of histone acetylation is reduced in the PDC mutants. Biochemical assay combined with Liquid Chromatography with mass spectrometry (LC-MS) demonstrates that PDC regulates the nuclear acetyl CoA concentration and ethylene induced nuclear translocated PDC is enzymatically active to generate acetyl CoA in the nucleus for histone acetylation, which is necessary for EIN2-C’s function in the histone acetylation regulation. Both genetics and molecular evidence show that EIN2-C is required for the nuclear translocation of PDC in the ethylene response. Altogether, this study reveals a previously unidentified mechanism by which the PDC complex is translocated to the nucleus where it interacts with EIN2-C to provide acetyl CoA for the elevation of histone acetylation at H3K14 and H3K23 to modulate the transcriptional regulation in the ethylene response.

## Results

### Nuclear localized pyruvate dehydrogenase complex interacts with EIN2-C

By tandem co-immunoprecipitation coupled with mass spectrometry (Co-IP/MS) using anti-EIN2-C native antibody and anti-HA antibody sequentially in the nuclear extracts from *EIN2-YFP-HA* plants treated with 4 hours of ethylene gas, three subunits from pyruvate dehydrogenase complex (PDC), E1-2, E2-2, and E3-2 (henceforth referred as E1, E2, and E3), were pulled down among the top hits (Fig. S1A-1C). The direct interactions between EIN2-C and these three PDC subunits identified in Co-IP/MS were confirmed by *in vitro* pull-down assays (Fig. 1A and Fig. S1D-F). In prokaryotes and eukaryotes, PDC consists of three catalytic subunits: a pyruvate dehydrogenase (E1), a dihydrolipoamide transacetylase (E2), and a dihydrolipoamide dehydrogenase (E3). This complex is canonically localized in the mitochondrial matrix where it catalyzes the conversion from pyruvate to acetyl CoA ^44^. The interaction between EIN2-C and PDC complex led us to examine whether it is possible that EIN2 is localized to the mitochondria. Co-localization assay of EIN2 with MitoTracker Red, a cell-permeant mitochondria-selective dye, showed that EIN2 and mitochondria were not co-localized either with or without the presence of ethylene (Fig. 1B). Given that PDC complex was pulled down from the nuclear fraction by EIN2-C Co-IP/MS, we then decided to confirm whether the interactions between EIN2-C and E1, E2, and E3 occur in the nucleus *in vivo*. We generated *EIN2-YFP-HA/E1-FLAG-BFP*, *EIN2-YFP-HA/E2-FLAG-BFP,* and *EIN2-YFP-HA/E3-FLAG-BFP* transgenic plants. By using EIN2-YFP-HA or PDC-FLAG-BFP as bait, we performed reciprocal *in vivo* co-immunoprecipitation (Co-IP) assays both in the nuclear fractions and in the cytosolic fractions from those 3-day-old etiolated transgenic seedlings treated with or without 4 hours of ethylene gas. The efficiency of nuclear-cytoplasmic fractionation procedure as well as the purity of cytosolic and nuclear fractions were assessed by multiple marker proteins (Fig. S1G). The interactions between EIN2-C and all three PDC subunits were detected only in the nuclear fractions with ethylene treatment, and no interactions were detected from cytosolic fractions regardless of the ethylene treatment (Fig. 1C-1E). Similar results were obtained by using the native EIN2-C antibody for the in vivo Co-IP assay in the *E1-YFP-HA*, *E2-FLAG-GFP*, and *E3-FLAG-GFP* transgenic plants (Fig. S1H-1J). To further evaluate the interaction between EIN2 and PDC in response to ethylene in the subcellular context, we conducted bimolecular fluorescence complementation (BiFC) assays by transient co-expression of EIN2-nYFP and E1-cYFP, E2-cYFP, and E3-cYFP, respectively, in tobacco epidermal cells. The result showed that EIN2-C physically interacts with PDC subunits only in the nucleus after ethylene treatment (Fig. 1F-1H). Furthermore, we examined the subcellular localization of EIN2-C and PDC subunits in response to ethylene in 3-day-old etiolated seedlings of *EIN2-YFP-HA/E1-FLAG-BFP*, *EIN2-YFP-HA/E2-FLAG-BFP,* and *EIN2-YFP-HA/E3-FLAG-BFP* plants. Clearly, EIN2-C was co-localized with E1, E2, and E3 in the nucleus respectively with the ethylene treatment (Fig. S1K-1P). Altogether, these results provide compelling evidence that PDC can accumulate in the nucleus to interact with EIN2-C in response to ethylene.

**Figure 1.**
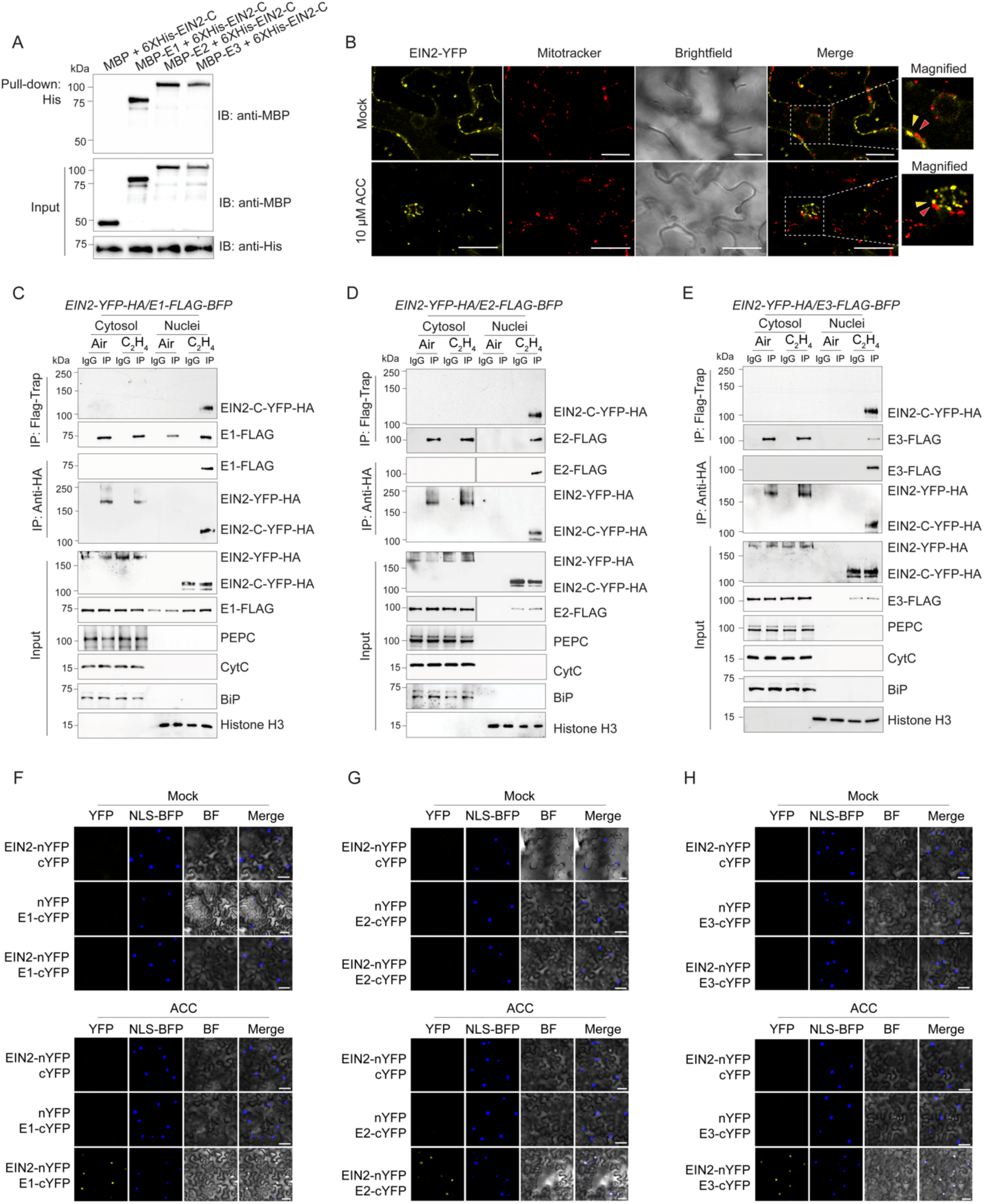
PDC interacts with EIN2-C in the nucleus in response to ethylene. (**A**) *In vitro* pull-down experiment to validate the interaction between EIN2-C and PDC E1, E2, and E3 subunits. (**B**) Subcellular localization of EIN2-YFP fusion protein in *Arabidopsis* leaf epidermal cells with MitoTracker Red staining without (upper panel) and with (lower panel) the presence of 10μM ACC. Red arrowhead indicates mitochondria signal by MitoTracker staining; yellow arrowhead indicates EIN2-YFP fluorescence signal. Scale bar is 20 μm. (**C**-**E**) *In vivo* co-immunoprecipitation assay to examine the interaction between EIN2-C and E1(**C**), E2 (**D**), and *E3* (**E**) in the indicated transgenic plants. 3-day-old etiolated seedlings carrying both PDC and EIN2 fusion proteins treated with air or 4 hours of ethylene gas were fractionated to isolate cytosol and nuclei for the immunoprecipitation with either Flag-Trap magnetic agarose (DYKDDDDK Fab-Trap) or anti-HA magnetic beads, respectively. The immunoprecipitation with IgG beads serves as a negative control. Phospho Enol Pyruvate Carboxylase (PEPC) was used as cytosolic marker protein; Cytochrome C (CytC) is a mitochondrial marker protein, BiP is an ER marker to assess nuclear extraction purity. Histone H3 is a loading control for nuclear fractions. (**F**-**H**) Confocal microscopy images of BiFC assay showing the interaction between EIN2 and PDC E1, E2, and E3 subunits in the nucleus after ethylene treatment. Agrobacteria containing indicated paired constructs was co-infiltrated into tobacco leaves and the YFP fluorescence was observed two days after infiltration with or without 4 hours of ACC treatment. NLS-BFP was used as nuclear marker. Scale bars is 50µm.

### Ethylene treatment induces the nuclear accumulation of PDC

We have observed the nuclear localization of PDC complex with the presence of ethylene (Fig. 1). To further confirm that the PDC nuclear translocation is induced by ethylene, we examined the nuclear appearance of E1, E2, and E3 proteins over a time series of ethylene gas treatments. E1 had a basal nuclear distribution in the absence of ethylene, but its nuclear levels were significantly elevated by ethylene treatment and were positively correlated with the duration of ethylene treatment while E1 cytoplasmic levels showed gradual decrease as the ethylene treatment prolongs (Fig. 2A, Fig. S2A, and Fig. S2B). Neither E2 nor E3 proteins were detected in the nucleus in the absence of ethylene, but these two proteins were accumulated in the nuclear fraction after 4 hours of ethylene treatment and their levels were also positively correlated with the duration of ethylene treatments (Fig. 2B, Fig. 2C, and Fig. S2C-2F). Similar to cytoplasmic E1, we also observed that cytoplasmic E2 and E3 protein levels decrease in response to ethylene (Fig. 2B, Fig. 2C, and Fig. S2C-2F). Notably, the total E1 and E2 subunit protein levels were not altered by the ethylene treatments but E3 total protein level showed slight elevation after 12h ethylene treatment. We then monitored the nuclear localization of PDC over a time series of ethylene treatments in living cells. In the absence of ethylene, a basal level of E1 was observed in the nucleus, but no nuclear E2 and E3 were observed (Fig. 2D-2F and Fig. S2G-2L). Upon ethylene treatment, E1, E2, and E3 accumulated in the nucleus, and their accumulation was positively associated with the duration of ethylene treatment (Fig. 2D-2F and Fig. S2G-2L). This provides an additional piece of cellular evidence that ethylene induces PDC nuclear accumulation.

**Figure 2.**
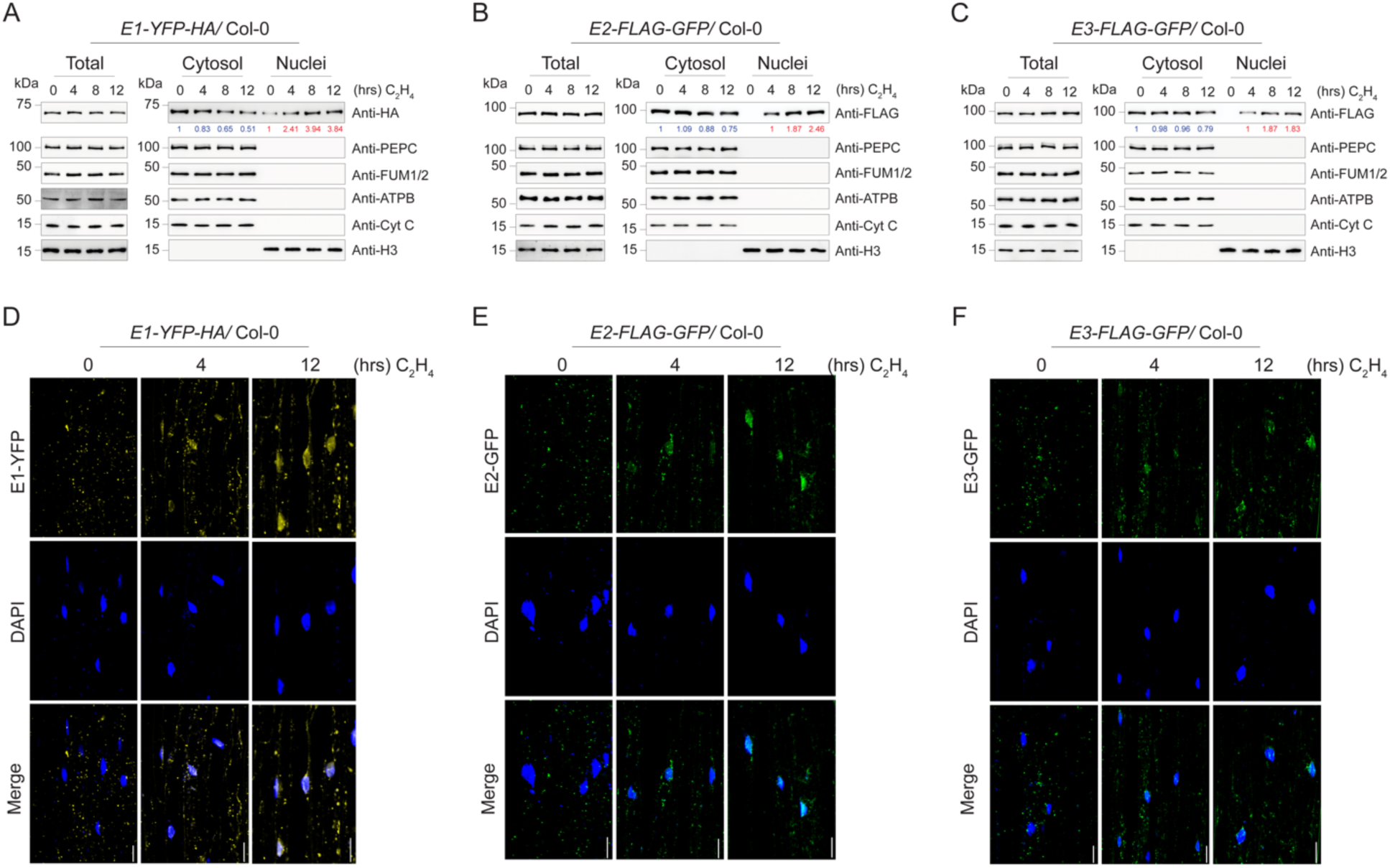
PDC accumulates in the nucleus in response to ethylene. (**A**-**C**) Fractionation western blot to examine the subcellular localization of PDC in of *E1-YFP-HA* (**A**), *E2-FLAG-GFP* (**B**), and *E3-FLAG-GFP* (**C**) transgenic plants with time series of ethylene gas treatments. PDC E1 was probed with anti-HA, and E2 and E3 were probed with anti-FLAG in total protein extracts, cytoplasmic fractions, and nuclear fractions. PEPC, ATPB, FUM1/2, Cyt C, and histone H3 were used to assess purities of nuclear and cytosolic fractionations and loading controls. Blue number indicates PDC band intensity that normalized to cytoplasmic PEPC signal. Red number indicates PDC band intensity that normalized to histone H3 signal. (**D**-**F**) Confocal microscopy images showing the subcellular localization of E1-YFP-HA (**D**), E2-FLAG-GFP (**E**), and E3-FLAG-GFP (**F**) with time series of ethylene gas treatments. DAPI staining labels nuclei. Scale bars is 20µm.

### PDC subunits are required for ethylene response

Given the interaction between PDC and EIN2-C occurs in the nucleus in response to ethylene, we decided to investigate the functions of PDC in the ethylene response. We first obtained two T-DNA insertion lines for *E1* (*e1-2-2* and *e1-2-4)* and two for *E3* (*e3-2-1* and *e3-2-2)*. RT-qPCR assays showed that the expression levels of these two genes were drastically reduced in their T-DNA insertion mutants (Fig. S3A-3D). No *E2* T-DNA homozygous plants were obtained due to homozygous lethality ^45^. Using CRISPR-Cas9 mutagenesis, we generated two weak alleles of *e2* mutants (*e2-2 #17* and *e2-2 #215*) that were variable (Fig. S3E-3G). Phenotypical assay showed that the *e2* single mutants, but not *e1* nor *e3* single mutant, displayed a mild ethylene insensitivity both in roots and in hypocotyls (Fig. 3A-3C). We then generated a variety of their higher order mutants and examined their ethylene responses (Fig. S3E-3G). The mild ethylene insensitive phenotype of the *e2* single mutant was significantly enhanced in *e1e2* (*e1-2-2 e2-2 #17* and *e1-2-4 e2-2 #215*) and *e2e3* (*e2-2 e3-2-1 #99* and *e2-2 e3-2-2 #67*) double mutants and in *e1e2e3* triple mutants (*e1-2-2 e2-2 e3-2-1 #167* and *e1-2-2 e2-2 e3-2-2 #67*) (Fig. 3D-3F). No significant difference was observed between the *e1e3* (*e1-2-2 e3-2-1* and *e1-2-2 e3-2-2*) double mutants and *e1* or *e3* single mutants (Fig. 3A-3F). The *e1e2*, *e2e3*, and *e1e2e3* ethylene insensitive phenotype were complemented by introducing *proE2:gE2-FLAG-GFP* into each mutant background, respectively (Fig. S3H and S3I). These genetic results demonstrated that PDC is involved in the ethylene response, E2 is the most important subunit genetically, and E1 and E3 enhance the function of E2 in the ethylene response.

**Figure 3.**
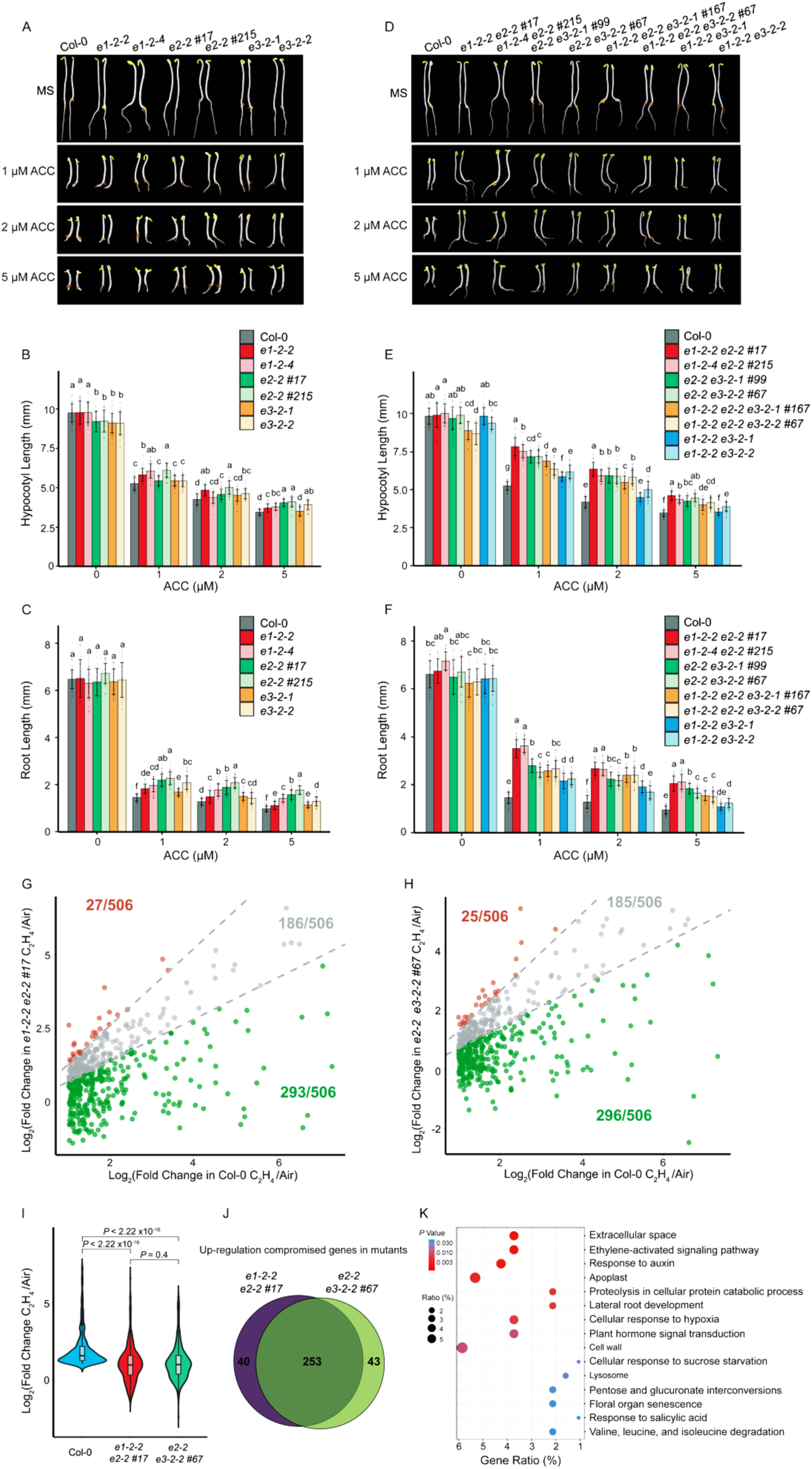
PDC is involved in the ethylene response. (**A**) Photographs of seedlings with mutations in individual PDC subunits and Col-0 grown on MS medium containing 1 μM, 2 μM, and 5 μM ACC or without ACC. (**B** and **C**) Measurements of hypocotyl lengths (**B**) and root lengths (**C**) of indicated mutants. Values are means ± SD of at least 30 seedlings. Different letters represent significant differences between each genotype calculated by a One-way ANOVA test followed by Tukey’s HSD test with *P* ≤0.05. (**D**) Photographs of representative double and triple PDC subunit mutants and Col-0 grown on MS medium containing 1 μM, 2 μM, and 5 μM ACC or without ACC were selected for the photograph. (**E** and **F**) Measurements of hypocotyl lengths and root lengths from indicated plants. Each value is means ± SD of at least 30 seedlings. Different letters indicate significant differences between different genotypes with *P* ≤0.05 that calculated by a One-way ANOVA test and followed by Tukey’s HSD test for multiple comparisons. (**G** and **H**) Scatter plots to compare the log_2_(Fold Change) of ethylene up-regulated genes in Col-0 with that in *e1-2-2 e2-2 #17* (**G**), or in *e2-2 e3-2-2 # 67* (**H**) in mRNA-seq. Each dot represents a gene that its transcription level is significantly elevated by ethylene gas treatment in Col-0 (log_2_(Fold Change) > 1, *p*-adjust < 0.05). (**I**) Violin plot of the distributions of log_2_(Fold Change) of ethylene up-regulated genes in Col-0 and in the indicated plants. *P* values were determined by a two-tailed *t* test. (**J**) Venn diagram to show the ethylene up-regulated genes that were compromised in *e1-2-2 e2-2 #17* and in *e2-2 e3-2-2 # 67*. (**K**) Gene ontology analysis of the genes that are transcriptionally co-compromised in both *e1-2-2 e2-2 #17* and in *e2-2 e3-2-2 #67*. Top 15 GO categories ranked by *P* values were plotted. Dot size represents the percentage of genes from each GO category of all co-compromised genes (Gene ratio %) and dot colors indicate the *P* value of the GO categories.

To further understand the functions of E1, E2, and E3 in the ethylene response, we generated their gain-of-function mutants and selected two individual transgenic lines of each with similar protein expression levels for further ethylene response analysis (Fig. S4A-4C). Compared to Col-0, all the plants that overexpressed *PDC E1, E2,* and *E3* subunits had enhanced ethylene responsive phenotype in the presence of ACC, but the phenotypes of *E2ox* and *E3ox* did not differ from Col-0 in the absence of ACC (Fig. S4D). This showed that the overexpression of *PDC E2* and *E3* alone was not sufficient to trigger ethylene response in the absence of the hormone when they were not accumulated in the nucleus. Interestingly, *PDC E1*ox displayed a mild ethylene hypersensitivity in the absence of ACC compared to Col-0, which is in line with the observation that there was a basal E1 in the nucleus without ethylene treatment (Fig. S4D). Next, we expressed *E1* and *E2* fused with nuclear localization signal peptide (NLS) derived from EIN2-C (*E1-NLS-GFP* and *E2-NLS-GFP*) in Col-0 (Fig. S4E). E1-NLS-GFP and E2-NLS-GFP fusion proteins could be localized to the nucleus in the absence of ethylene; more importantly, the *E1-NLS-GFP* and *E2-NLS-GFP* plants displayed a clear ethylene hypersensitivity phenotype even in the absence of ethylene (Fig. S4F and S4G). Together, these data demonstrate that the nuclear localization of PDC can trigger the ethylene response.

In order to evaluate the function of PDC in ethylene response at a molecular level, we performed RNA sequencing (mRNA-seq) using *e1-2-2 e2-2 #17* and *e2-2 e3-2-2 #67* double mutants treated with or without 4 hours of ethylene gas (Fig. S5). We found that about 50% genes up-regulated by ethylene in Col-0 were not differentially expressed in either of the two mutants in response to ethylene (Fig. S6A and S6B). For further statistical quantification, we plotted the log_2_ fold change (log_2_FC) of each gene up-regulated by ethylene in Col-0 against that in each double mutant. These genes were then divided into three groups: elevated group (log_2_FC in mutants is 30% greater than in Col-0), unchanged group (log_2_FC in mutants is within 30% of that in Col-0), and compromised group (log_2_FC in mutants is 30% less than that in Col-0). We found that the elevation of more than half of ethylene-up regulated genes in Col-0 was significantly compromised in both double mutants (Fig. 3G-3I). Importantly, most of the compromised genes identified from *e1-2-2 e2-2 #17* seedlings were also compromised in the *e2-2 e3-2-2 #67* seedlings (Fig. 3J), and the log_2_FC profile showed a high similarity (Fig. 3I and Fig. S6C). Further gene ontology analysis using those upregulation compromised genes showed that the ethylene signaling genes were overrepresented (Fig. 3K and Fig. S6D). Together, these data provide genetic and molecular evidence that E1, E2, and E3 function coordinately to regulate ethylene response.

### PDC regulates ethylene-dependent elevation of histone acetylation at H3K14 and H3K23

PDC catalyzes the synthesis of acetyl CoA, the substrate of histone acetylation. Given that the acetylation of histone at H3K14 and H3K23 is induced by ethylene, we compared their global levels in Col-0 and in different *pdc* mutants. The ethylene-induced global elevation of H3K14ac and H3K23ac in Col-0 was still detected in *e1-2-2 e2-2 #17* but with lower levels. Whereas, ethylene-induced global elevation was not detected in *e2-2 e3-2-2 #67*, nor in *e1-2-2 e2-2 e3-2-2 #67* (Fig. 4A). In contrast, the H3K14ac and H3K23ac levels were elevated in the plants that overexpress *E2* with ethylene treatment, and in the *E2-NLS-GFP* transgenic plants even without ethylene treatment (Fig. 4B). We then conducted H3K14ac and H3K23ac chromatin immunoprecipitation sequencing (ChIP-seq) in *e1-2-2 e2-2 #17* (Fig. S7A-7D). H3K14ac and H3K23ac levels were reduced in *e1-2-2 e2-2 #17* compared to that in Col-0 over genes that were transcriptionally compromised in both *e1-2-2 e2-2 #17* and *e2-2 e3-2-2 #67* plants, and the ethylene-induced elevation was significantly impaired (Fig. 4C, 4D, and Fig. S7E-7H). When the ChIP signals within the first 500bp downstream of transcription starts sites was evaluated, an even more significant reduction was detected in *e1-2-2 e2-2 #17* both with and without ethylene treatments (Fig. 4E and 4F). We then conducted H3K14ac and H3K23ac ChIP-qPCR assays in the selected targets, and the results further validated the ChIP-seq data (Fig. S7I and S7J). Given that *e2-2 e3-2-2 #67* and *e1-2-2 e2-2 e3-2-2 #67* has a similar ethylene responsive phenotype as *e1-2-2 e2-2 #17*, we also conducted ChIP-qPCR assays in those two mutants, and a similar result was obtained (Fig. S7I and S7J). Altogether, these results suggest that PDC regulates the histone acetylation H3K14ac and H3K23ac in response to ethylene.

**Figure 4.**
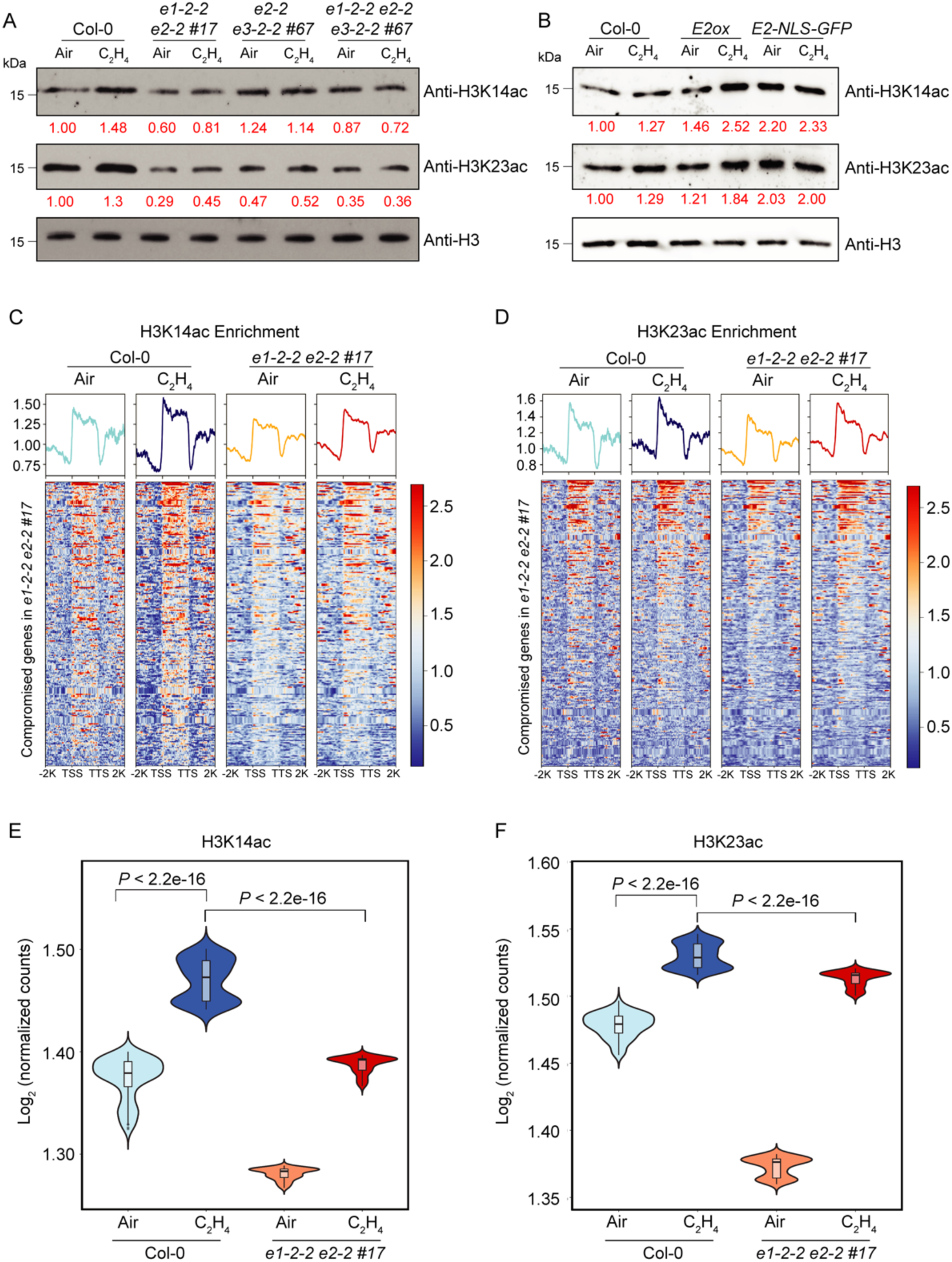
PDC is involved in the ethylene-induced histone acetylation. (**A**) Western blot analysis of total histone acetylation of H3K14 and H3K23 in response to ethylene in different genetic backgrounds as indicated in the figure. Anti-H3 western blot served as a loading control. (**B**) H3K14ac and H3K23ac levels in Col-0, *E2ox,* and *E2-NLS-GFP* treated with air or 4 hours of ethylene gas. Histone H3 served as a loading control. Red number indicates the quantification of H3K14ac and H3K23ac western blot signal intensity normalized with that of histone H3. (**C** and **D**) Heatmaps of H3K14ac (**C**) and H3K23ac (**D**) ChIP-seq signal (log_2_ ChIP signal) from the genes that their ethylene-induced expressions are co-compromised in *e1-2-2 e2-#17* and *e2-2 e3-2-2 #67*. TSS, transcription start site; TTS, transcription termination site. (**E** and **F**) Violin plots illustrating log_2_ normalized H3K14ac ChIP signal (**E**) and H3K23ac ChIP signal (**F**) from 500bp downstream of TSS that in the genes that were co-compromised in the *pdc* double mutants in the indicated genotypes and conditions. *P* values were calculated by a two-tailed *t* test.

### Enzymatically active nuclear PDC regulates acetyl CoA production in the nucleus

Because the biological function of PDC is to generate acetyl CoA, we hypothesized that the reduced histone acetylation level in the plants that lack PDC subunits results from the decreased acetyl CoA in the nucleus. We first measured the nuclear acetyl CoA concentrations in Col-0 and in the *e1-2-2 e2-2 e3-2-2 #67* triple mutant treated with 4 hours of ethylene gas by liquid chromatography–(LC) coupled to MS (LC–MS), and the levels of nuclear acetyl CoA were significantly decreased in the *pdc* triple mutant compared to levels in Col-0 (Fig. 5A and 5B). We then decided to examine whether the nuclear PDC was functional to synthesize acetyl CoA. It has been shown that dephosphorylated E1 is required for the PDC activity ^44,46^, therefore, we first examined the phosphorylation status of E1 in the nucleus in response to ethylene. Western blot assay by anti-pSer antibody showed that most E1 was in a dephosphorylated state in the nucleus after ethylene treatment (Fig. 5C). Phos-tag electrophoresis assay confirmed this result (Fig. 5D). Next, we conducted LC-MS/MS analysis of E1 phosphorylation. We detected phosphorylation on S292 (Fig. S8A), a residue that is evolutionary conserved with the reported inhibitory phosphorylation site in human E1 (Fig. S8B)^46^. Further quantification of E1 phosphopeptides by MS showed that the ratio of phosphorylated E1 to non-phosphorylated E1 in the nucleus was significantly lower than that in the cytosol after 4 hours of ethylene treatment (Fig. 5E), suggesting that the dephosphorylated E1 is the main species in the nucleus to function in the presence of ethylene. Next, we assessed E1 activities in the nucleus using a pyruvate dehydrogenase (PDH) activity assay. E1 activity was detected in the nucleus in Col-0 and its activity was elevated by the ethylene treatment (Fig. 5F). But the ethylene-induced E1 activity was not detected in *e1-2-2 e2-2#17* or in *e1-2-2 e2-2 e3-2-2#67* (Fig. 5F and Fig. S8C). To further directly monitor PDC acetyl CoA biosynthesis enzyme activity in the nucleus in response to ethylene, we employed a ^13^C isotopic tracing experiment. By feeding 2,3-^13^C_2_ pyruvate to the nuclear extracts and followed by the LC-MS/MS, we are able to measure 1,2-^13^C_2_ labeled-acetyl CoA (1,2-^13^C_2_ acetyl CoA) that will be converted from 2,3-^13^C_2_ pyruvate by PDC complex (Fig. 5G and Fig. S8D). This experiment assesses the nuclear PDC activity since PDC is the only enzyme that synthesizes acetyl CoA from pyruvate. As shown in Fig. 5H, we detected 1,2-^13^C_2_ acetyl CoA in the nuclear extracts of Col-0 treated with 4 hours of ethylene gas, and the 1,2-^13^C_2_ acetyl CoA levels were significantly increased in the presence of ethylene (Fig. 5H and Fig. S8E). This suggests that functional PDC complex is accumulated in the nucleus in response to ethylene in Col-0. Further comparison showed no significant differences in the levels of 1,2-^13^C_2_ acetyl CoA in the nuclei of Col-0 compared to that of *e1-2-2 e2-2 #17* or of *e1-2-2 e2-2 e3-2-2 #*67 in the absence of ethylene (Fig. 5H and Fig. S8E). However, the ethylene-induced elevation of 1,2-^13^C_2_ acetyl CoA detected from Col-0 nucleus was significantly reduced from the nucleus of *e1-2-2 e2-2 #17* or *e1-2-2 e2-2 e3-2-2#67* (Fig. 5H), providing further biochemical evidence that functional PDC is translocated into the nucleus in response to ethylene to synthesize acetyl CoA.

**Figure 5.**
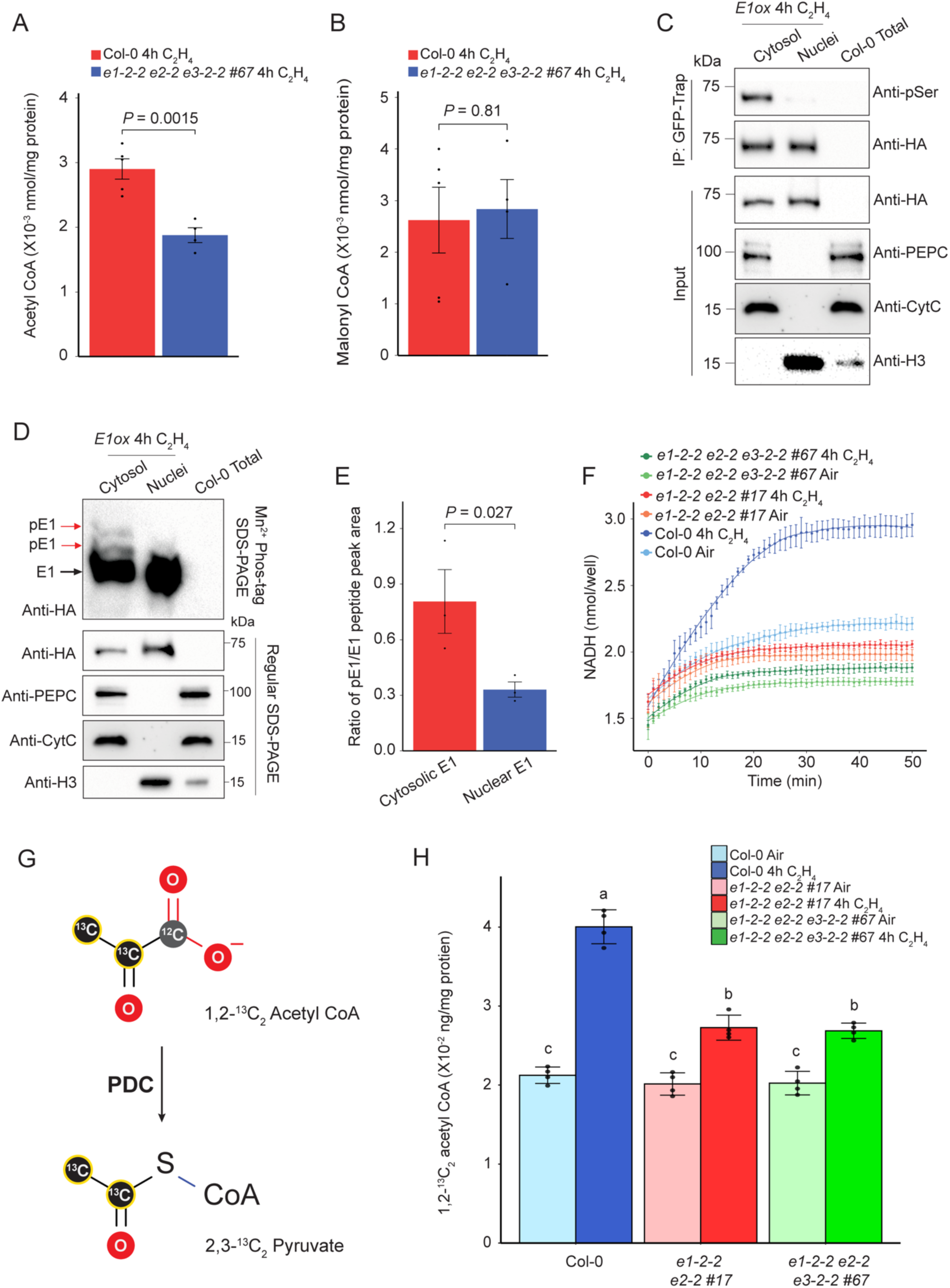
Nuclear PDC is enzymatically active to produce acetyl CoA. (**A** and **B**) LC-MS detection of acetyl CoA (**A**) and malonyl CoA (**B**) in purified nuclei from Col-0 and *e1-2-2 e2-2 e3-2-2 #67* etiolated seedlings treated with 4 hours of ethylene gas. Total protein mass was used to normalize metabolite concentration. *P* values were calculated by a two-tailed *t* test and each data point was plotted as a dot in the bar graph. (**C**) Western blot analysis of the phosphorylation status of E1 using anti-pSer antibody in the cytosolic fraction and nuclear fractions from *E1ox* seedlings treated with 4 hours of ethylene gas. (**D**) Phos-tag phosphoprotein mobility shift gel of E1 phosphorylation status in the cytosolic fraction and nuclear fractions of *E1ox* etiolated seedlings treated with 4 hours of ethylene gas. Black arrow shows non-phosphorylated E1; red arrow indicates phosphorylated E1 protein. PEPC, CytC, and H3 were used to assess purities of nuclear and cytosolic fractionations. (**E**) The ratios of phosphorylated to non-phosphorylated YHGHpSMSDPGSTYR E1 peptides in the cytosolic versus in the nuclear fractions from *E1ox* with 4 hours of ethylene gas treatment. Peak area values were collected from three replicates. *P* value was calculated by *t*-test. (**F**) Pyruvate dehydrogenase activity assays in the nuclear fraction from Col-0 and indicated *pdc* mutants treated with 4 hours of ethylene gas. (**G**) Schematic diagrams to show the conversion of 2,3-^13^C_2_ pyruvate to 1,2-^13^C_2_ acetyl CoA. (**H**) Normalized LC MS/MS measured the concentration of 1,2-^13^C_2_ acetyl CoA that converted from 2,3-^13^C_2_ pyruvate by using the nuclear extracts from indicated plants with or without 4 hours of ethylene gas treatment. Quantification from four replicates were normalized by total protein input. Different letters represent significant differences between each group calculated by a one-way ANOVA test followed by Tukey’s HSD test.

### EIN2 is required for PDC nuclear translocation and function in response to ethylene

Given the fact that EIN2-C interacts with PDC in the nucleus, we next explored the genetic relationship between EIN2-C and PDC subunits. By introducing *E1ox, E2ox*, and *E3ox* into the *ein2-5* null mutant, we obtained *E1ox/ein2-5, E2ox/ein2-5*, and *E3ox/ein2-5* plants with comparable protein expression levels in *E1ox/*Col-0*, E2ox/* Col-0, and *E3ox/* Col-0 separately (Fig. S9A-9C). We then compared their ethylene response with their parental plants and Col-0. The ethylene hypersensitivity induced by PDC overexpression was entirely abolished in all *E1ox/ein2-5, E2ox/ein2-5*, and *E3ox/ein2-5* plants; these plants displayed the same complete ethylene insensitivity as *ein2-5* mutant (Fig. 6A). Similarly, the ethylene hypersensitivity induced by PDC overexpression was entirely abolished in all *E1ox/ein3-1eil1-1, E2ox/ein3-1eil1-1*, and *E3ox/ein3-1eil1-1* plants (Fig. S9A-9D). These two genetic results suggest that both EIN2 and EIN3/EIL1 are required for the function of PDC in the ethylene response.

**Figure 6.**
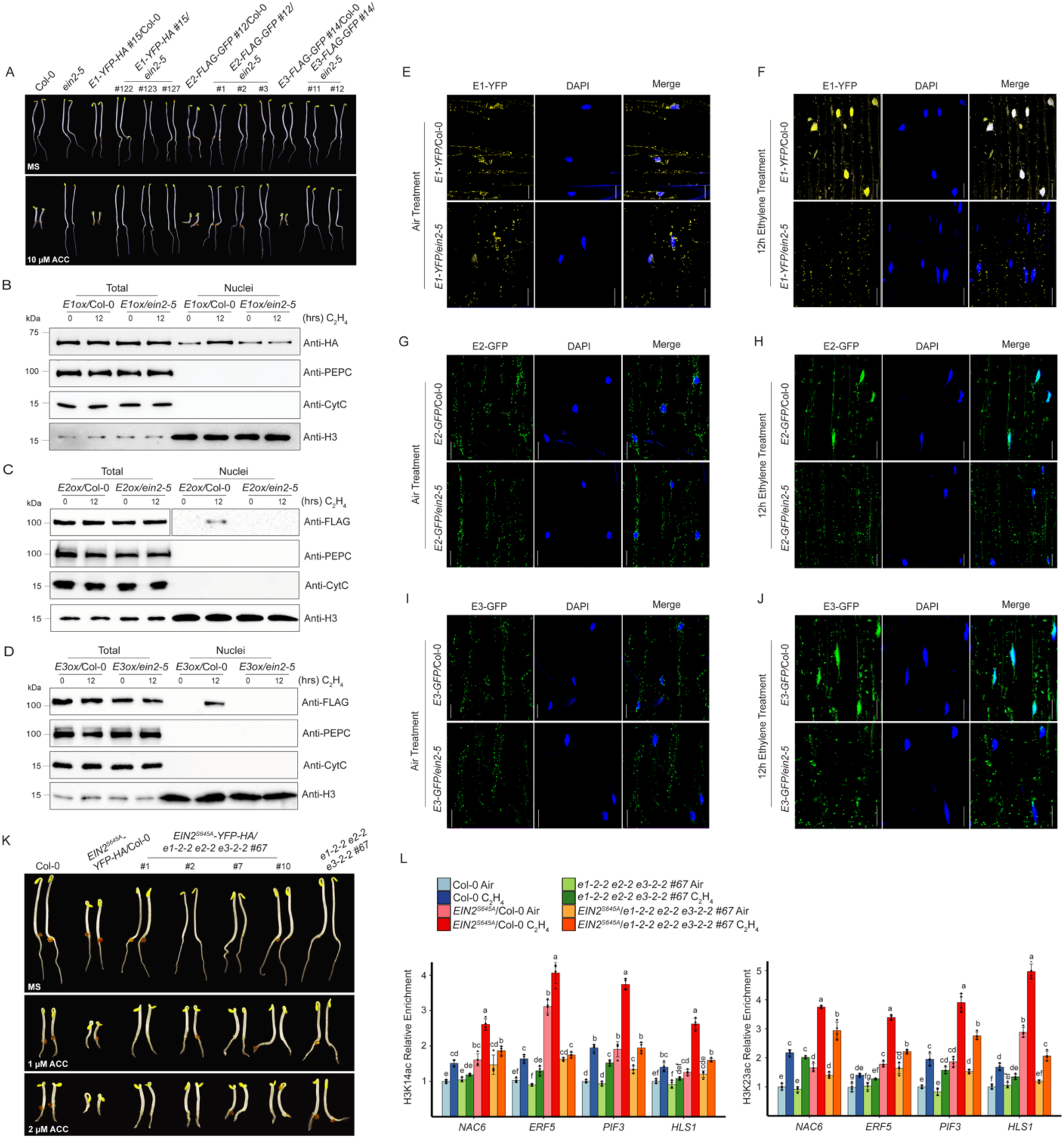
PDC functions in the ethylene response in an EIN2-C-dependent manner. (**A**) Epistasis analysis of PDC *E1ox*, *E2ox*, and *E3ox* in the *ein2-5* mutant. The seedlings were grown on MS medium containing with or without 10μM ACC in the dark for 3 days before being photographed. (**B**-**D**) Examination of the protein levels of E1 in *E1ox/*Col-0 and *E1ox/ein2-5* (**B**), E2 in *E2ox/*Col-0 and *E2ox/ein2-5* (**C**), and E3 in *E3ox/*Col-0 and *E3ox/ein2-5* (**D**) with 0 or 12-hour ethylene gas treatment. PEPC is cytoplasmic marker; CytC is a mitochondrial marker; histone H3 is a nuclear marker for a loading control. (**E**-**J**) Subcellular localization of E1-YFP-HA (**E** and **F**), E2-FLAG-GFP (**G** and **H**), and E3-FLAG-GFP (**I** and **J**) in Col-0 or *ein2-5* mutant without or with 12 hours of ethylene gas treatment, respectively. Scale bars is 20µm. (**K**) Photographs of 3-day old etiolated *EIN2^S^*^645^*^A^/*Col-0 and *EIN2^S^*^645^*^A^/e1e2e3* seedlings grown on MS medium containing indicated ACC in the dark for 3 days before being photographed. (**L**) ChIP-qPCR to evaluate H3K14ac (upper panel) and H3K23ac (lower panel) enrichment over selected genes in the indicated etiolated seedlings treated with air or 4 hours of ethylene gas. Different letters represent significant differences between each genotype and treatment condition that are calculated by a one-way ANOVA test followed by Tukey’s HSD test with *P* ≤0.05.

To further confirm the EIN2 dependency of PDC in ethylene response, we examined the cellular localization of PDC in response to ethylene in the *ein2-5* mutant. We found that the ethylene-induced nuclear accumulation of E1, E2, and E3 was abolished in *E1ox/ein2-5, E2ox/ein2-5*, and *E3ox/ein2-5* (Fig. 6B-6J). Similarly, no obvious ethylene-induced nuclear accumulation of PDC proteins was observed in *E1ox/ein3-1 eil1-1, E2ox/ein3-1 eil1-1*, and *E3ox/ein3-1 eil1-1* (Fig. S9E-9J). ChIP-qPCR analyses in selected ethylene-responsive target genes (Fig. S7E-7H) showed that the elevation of H3K14ac and H3K23ac levels detected in *E2ox* were not detected in the *E2ox/ein2-5* plants, and the levels were similar to those in the *ein2-5* mutant both with and without ethylene treatments (Fig. S9K). In addition, the enhanced transcriptional activation in response to ethylene in *E2ox* was not detected in *E2ox/ein2-5* plants (Fig. S9L). Thus, our genetic and molecular evidence supports the conclusion that PDC functions in the ethylene response in an EIN2 dependent manner.

Acetyl CoA is important for histone acetylation and EIN2-C is essential for the ethylene-induced histone acetylation elevation; therefore, we further investigated whether PDC regulates the function of EIN2-C. We crossed *EIN2^S^*^645^*^A^*, in which a point mutation at Ser645 mimics EIN2 constitutive dephosphorylation resulting in EIN2-C cleavage and nuclear accumulation, into *e1-2-2 e2-2 e3-2-2 #67* (*e1e2e3*) background to generate *EIN2^S^*^645^*^A^/e1e2e3* plants (Fig. S10A). Phenotypic analysis of *EIN2^S^*^645^*^A^/*Col-0 and *EIN2^S^*^645^*^A^/e1e2e3* showed that the ethylene hypersensitivity caused by *EIN2^S^*^645^*^A^*was partially rescued in the *e1e2e3* mutant (Fig. 6H). But the nuclear accumulation of EIN2-C was comparable in the *EIN2^S^*^645^*^A^/*Col-0 and in *EIN2^S^*^645^*^A^/e1e2e3* (Fig. S10B and S10C), showing that PDC participates in the downstream ethylene signaling and cellular response mediated by EIN2-C rather than the initial steps of EIN2-C cleavage and translocation into the nucleus in ethylene signaling pathway. We then compared the H3K14ac and H3K23ac levels at selected ethylene-responsive genes in *EIN2^S^*^645^*^A^/*Col-0 and *EIN2^S^*^645^*^A^/e1e2e3* plants by ChIP-qPCR. We found that the enhanced H3K14ac and H3K23ac in *EIN2^S^*^645^*^A^/*Col-0 was reduced in *EIN2^S^*^645^*^A^/e1e2e3,* although levels were still higher than that in Col-0 (Fig. 6I). Consistently, the enhancement of target gene expression in *EIN2^S^*^645^*^A^/*Col-0 was also partially rescued in *EIN2^S^*^645^*^A^/e1e2e3* (Fig. S10D). These findings suggest that PDC in the nucleus produces acetyl CoA that is necessary for EIN2-C-regulated histone acetylation to mediate the ethylene response.

## Discussion

This study reveals a novel molecular mechanism by which mitochondria resided PDC complex translocates into the nucleus to provide acetyl CoA necessary for EIN2-C mediated histone acetylation and subsequent transcription activation in response to ethylene (Fig. S11). In the presence of ethylene, PDC subunits E1, E2, and E3 accumulate in the nucleus to provide acetyl CoA for EIN2 mediated histone acetylation elevation at H3K14 and H3K23 through their interaction with EIN2-C, initiating the downstream ethylene-induced transcriptional cascade (Fig. S11). Our research establishes a direct link between cell metabolism and histone modification in the ethylene response that is mediated by the key factor EIN2, opening a new avenue for the study of how plant hormone and metabolisms function to ensure the chromatin and transcription regulation.

Cellular metabolism is a series of important biochemical reactions fueling development with energy and biomass, and chromatin is a mighty consumer of cellular energy generated by metabolism. The balance between metabolism and chromatin activities ensures cell homeostasis and normal cell growth. PDC functioning as a partner of EIN2-C to regulate histone acetylation levels at H3K14 and H3K23 marks in response to ethylene treatment suggests that EIN2-C recruits translocated PDC to the ethylene-responsive gene loci to provide a sufficient level of local acetyl CoA to achieve histone acetylation at H3K14 and H3K23 by HATs. Production of acetyl CoA within the nucleus may promote its availability to histone acetyltransferase to facilitate a rapid ethylene response. Thus, identification of the histone acetyl transferases that function together with EIN2-C and PDC complex will be an immediate goal.

In the nucleus, acetyl CoA production machineries are finely tuned to control the local metabolite levels at certain gene loci to regulate chromatin modification by interacting with different partners. For instance, PDC E2 subunit has been reported to function in the same complex with pyruvate kinase M2 (PKM2), histone acetyltransferase p300, and the transcription factor AhR to regulate histone acetylation for transcription regulation to facilitate cell proliferation^47^. Nuclear metabolic enzyme Acetyl CoA Synthetase 2 (ACSS2) associates with the histone acetyltransferase CREP binding protein (CBP) to regulate the expression of key neuronal genes through histone acetylation in the mouse hippocampus^48^. In response to glucose deprivation, ACSS2 can also constitute a complex with the transcription factor EB to activate lysosomal and autophagosomal genes after its nuclear translocation^42^. In ethylene response, EIN3 is the key transcription factor that determines the target gene for transcription regulation^25,27^; thus, it will be interesting to explore the connection between EIN3 and PDC complex.

In mammal, three main cytosolic enzymes that synthesize acetyl CoA, including PDC, Acyl-CoA Synthetase Short Chain Family Member (ACSS2) and ATP Citrate Lyase (ACLY or ACL), were shown to localize to the nucleus for different biological functions^38,40,42,43,47–49^. PDC complex was first reported to localize to the nucleus in cell division and synthesize acetyl CoA to regulate cell proliferation through histone acetylation^40^. ACSS2 catalyzes the conversion of acetyl CoA from acetate in the nucleus for the regulation of long-term spatial memory and glucose starvation response^42,48^; the nuclear ACLY utilizes citrate to produce acetyl CoA to promote histone acetylation at double strand break sites for DNA repair^38^. Recently, ACL subunit A2 (ACLA2) has been reported to function with HAT1 in rice to regulate cell division in developing endosperm, suggesting the evolutionarily conserved mechanism of metabolic regulation on epigenetic modification in both plant and mammal species^50^. We have noticed that although the ethylene induced elevation of H3K14ac and H3K23ac is compromised in the *pdc* mutant, some extent of ethylene response still remains. It is possible that since we could not obtain the *e2* null mutant because of its lethality, the weak alleles we obtained still have partially functional E2 and therefore manifest a rather weakened ethylene response. However, we could not exclude the possibility that the other acetyl CoA producing enzymes will be functioning in the nucleus to modulate histone acetylation in response to ethylene in the absence of the nuclear PDC in *pdc* mutants. Investigating the involvement of other acetyl CoA synthesis pathways, such as ACSS^42,48^ and ACLY^38^, in the ethylene response will potentially address the question.

In this research, we found that ethylene treatment triggers the translocation of PDC from the cytosol to the nucleus to facilitate H3K14ac and H3K23ac elevation for ethylene responsive transcriptional regulation. However, the question remains how the translocation of PDC complex occurs at molecular level because PDC is a large protein complex without known nuclear localization signal (NLS). Recent studies have shown that specialized contact points between mitochondria and nucleus aid in the exchanges of metabolites and proteins between these two organelles, and more importantly, mammalian PDC directly enters the nucleus at those contact points across the nuclear envelop through mitofusin 2 (MFN2)-mediated mitochondria-nucleus tethering, which is independent from the nuclear pore complex (NPC)^51,52^. However, whether it is possible that nuclear PDC proteins are transported into the nucleus from the cytosol following their protein synthesis is unknown. Investigating how *Arabidopsis* PDC enters the nucleus and whether its nuclear entry follows the similar NPC-independent mechanism to regulate histone acetylation in response to ethylene will provide more insights into the mitochondria and nucleus retro-communication in the establishment of the proper ethylene response, providing a new perspective of metabolic regulation in plant hormone response research.

## Acknowledgements

We thank D. Hernandez for lab maintenance. We thank Dr. C. Petucci and Metabolomics Core of Cardiovascular Institute at University of Pennsylvania for nuclear acetyl CoA LC/MS quantification and Dr. K.S. Browning for nuclear-cytoplasmic fractionation setup. We thank P. Oliphint, A. Webb, and Microscopy and Imaging Facility at The University of Texas at Austin for the support of confocal microscope imaging. E1 phosphorylation MS analysis was supported by the Carnegie endowment fund to the Carnegie mass spectrometry facility, and we thank Andres V. Reyes for his assistance in MS analysis. We also thank *Arabidopsis* Biological Resource Center for the *Arabidopsis* T-DNA insertion mutants. Z.S. was supported by the Research Awards from Department of Integrative Biology and the Graduate Continuing Fellowship from The University of Texas at Austin. This work was supported by grants from the National Institute of Health to H.Q. (NIH-2R01 GM115879).

## Author contributions

Z.S. and H.Q. designed the study. Z.S. performed most of the experiments and analysis. L.B. performed LC-MS/MS analysis of 1,2-^13^C_2_ acetyl CoA and Y.B. performed MS analysis of E1 phosphorylation. Z. Shen performed Co-IP/MS. J.G.B. and P.K. helped genetic material generation and imaging. S.K.A., T.J.D., and M.A.B. helped biochemical assays and high-throughput sequencing and contributed to manuscript editing. S.-L.X., Z.-Y.W., S.P.B. and H.Q. supervised the study. Z.S. and H.Q. wrote the paper.

## Data and materials availability

Further information and requests for all unique materials generated in this study may be directed to and will be fulfilled by the corresponding author Hong Qiao. The high-throughput sequencing data generated in this study have been deposited in the Gene Expression Omnibus (GEO) database (accession no. GSE212540).

