## Supplemental Figures and Tables for "Nuclear Pyruvate Dehydrogenase Complex Regulates Histone Acetylation and Transcriptional Regulation in the Ethylene Response"

### Supplemental Materials

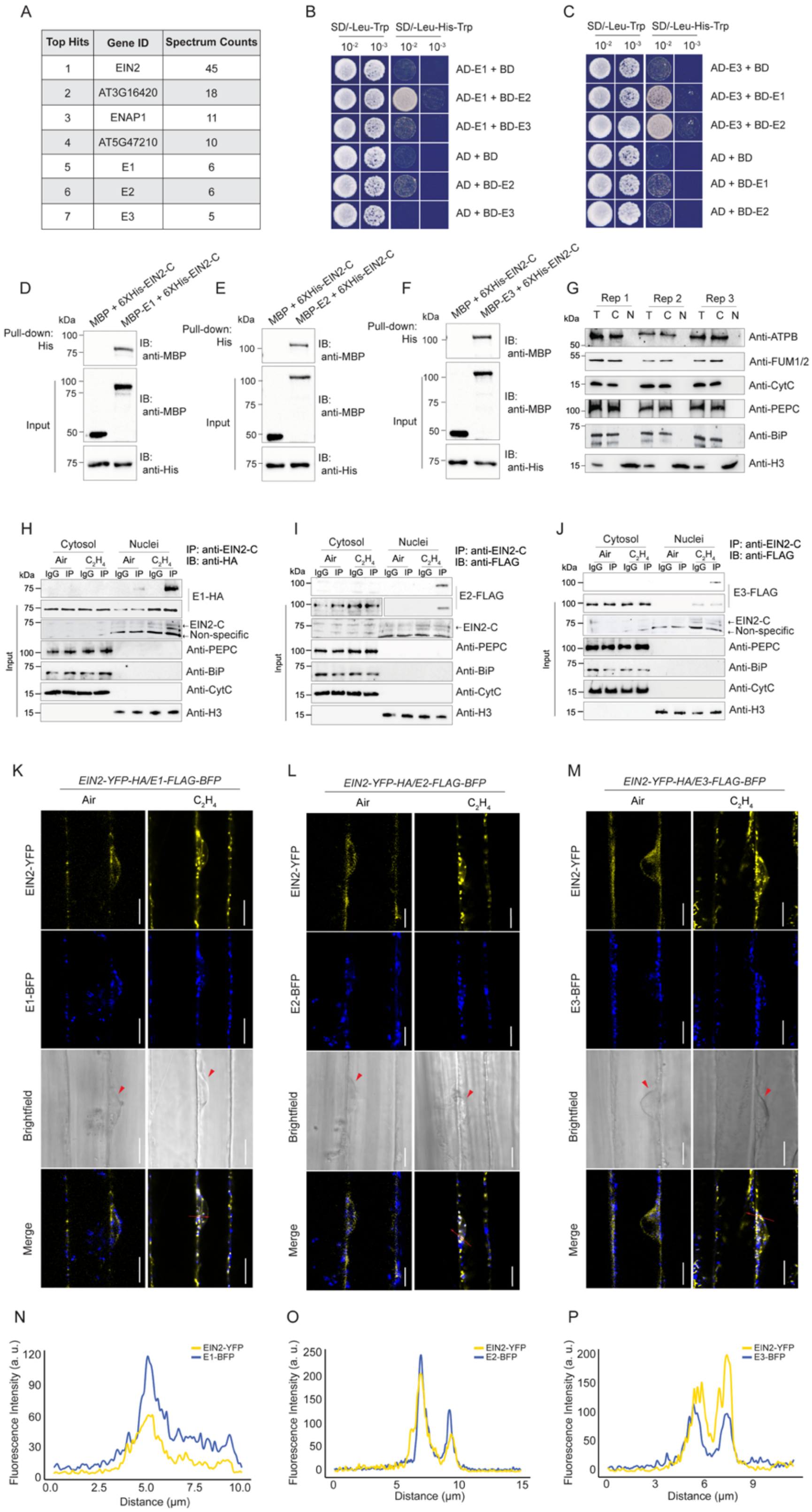

**Supplemental Figure 1. Related to Figure 1. Physical interactions between PDC E1, E2, and E3 and between EIN2-C and PDC subunits and nuclear co-localizations between EIN2-C and PDC subunits with ethylene treatment.**

(**A**) IP-MS spectral counts of part of EIN2-C interacting proteins with 4 hours of ethylene gas treatment. (**B** and **C**) Yeast two-hybrid assays showing that E1, E2, and E3 interact with each other to form a protein complex. BD: GAL4 DNA binding domain; AD: GAL4 activation domain. E1 was fused to AD and used as bait proteins in (**B**) and AD-E3 was used as bait proteins in (**C**). Left panels: Yeasts grown on two-dropout medium (SD/-Leu-Trp) served as a loading control; Right panels: Yeast grown on selective three-dropout medium (SD/-Leu-His-Trp). (**D**-**F**) Pull-down assay to examine the interactions between EIN2-C and PDC E1 (**D**), E2 (**E**), and E3 (**F**), respectively. His-tagged EIN2-C and MBP-tagged PDC subunits were purified from *E. coli* and subject to pull-down assays. MBP was used as a negative control. (**G**) Assessment of the purities of cytosolic and nuclear fractions. Three-day-old etiolated seedlings treated with 4 hours ethylene were applied to the nuclear-cytoplasmic fractionation procedure as described in the Materials and Methods section. Three independent biological replicates were included. Anti-ATP synthase Beta (anti-ATPB), anti-Fumarase 1+2 (anti-FUM1/2), and anti-Cytochrome C (anti-Cyt C) were used to evaluate the presence of mitochondrial proteins. Anti- PEP Carboxylase (anti-PEPC) was used to examine the cytoplasmic fraction and anti-Lumenal-binding protein (anti-BiP) serves as ER control. Anti-Histone H3 (anti-H3) served as nuclear protein loading. Rep 1, Rep 2, and Rep 3 represent replicate 1, replicate 2, and replicate 3, respectively. T represents total extract, C represents cytosolic fraction, and N represents nuclear fraction. (**H**-**J**) *In vivo* co-immunoprecipitation experiments to confirm the interaction between EIN2-C and PDC. Cytosolic and nuclear fractions from 3-day-old etiolated seedlings of *35S:E1-YFP-HA* (**H**), *35S:E2-FLAG-GFP* (**I**), and *35S:E3-FLAG-GFP* (**J**) treated with or without 4 hours of ethylene gas were used for the immunoprecipitation with anti-EIN2-C antibody. The immunoprecipitation with IgG beads was used as a negative control. E2 from the nucleus was detected in the IP products in the presence of 4 hours of ethylene gas (**I**). (**K**-**M**) Confocal microscopy images showing the co-localization of EIN2-C-YFP with PDC subunits E1-, E2- and E3-BFP in response to ethylene. The images were collected from the hypocotyls of 3-day old etiolated seedlings as indicated in the figure with or without 4 hours of ethylene treatment. Red arrowhead indicates nuclei in the brightfield. Red arrow represents the transection used for measuring fluorescence intensity as indicated in the figure. Scale bars is 10μm. (**N**-**P**) Fluorescence intensity measurements of confocal images from Fig. 1F-1H (lowest panel under ethylene treatment) to show co-localization of EIN2-C with PDC subunits in the nucleus with ethylene treatment. Yellow line indicates EIN2-YFP fluorescence intensity and blue line indicates PDC E1-BFP (**N**), E2-BFP (**O**), or E3-BFP (**P**) fluorescence intensity, respectively. Scale bars: 10μm.

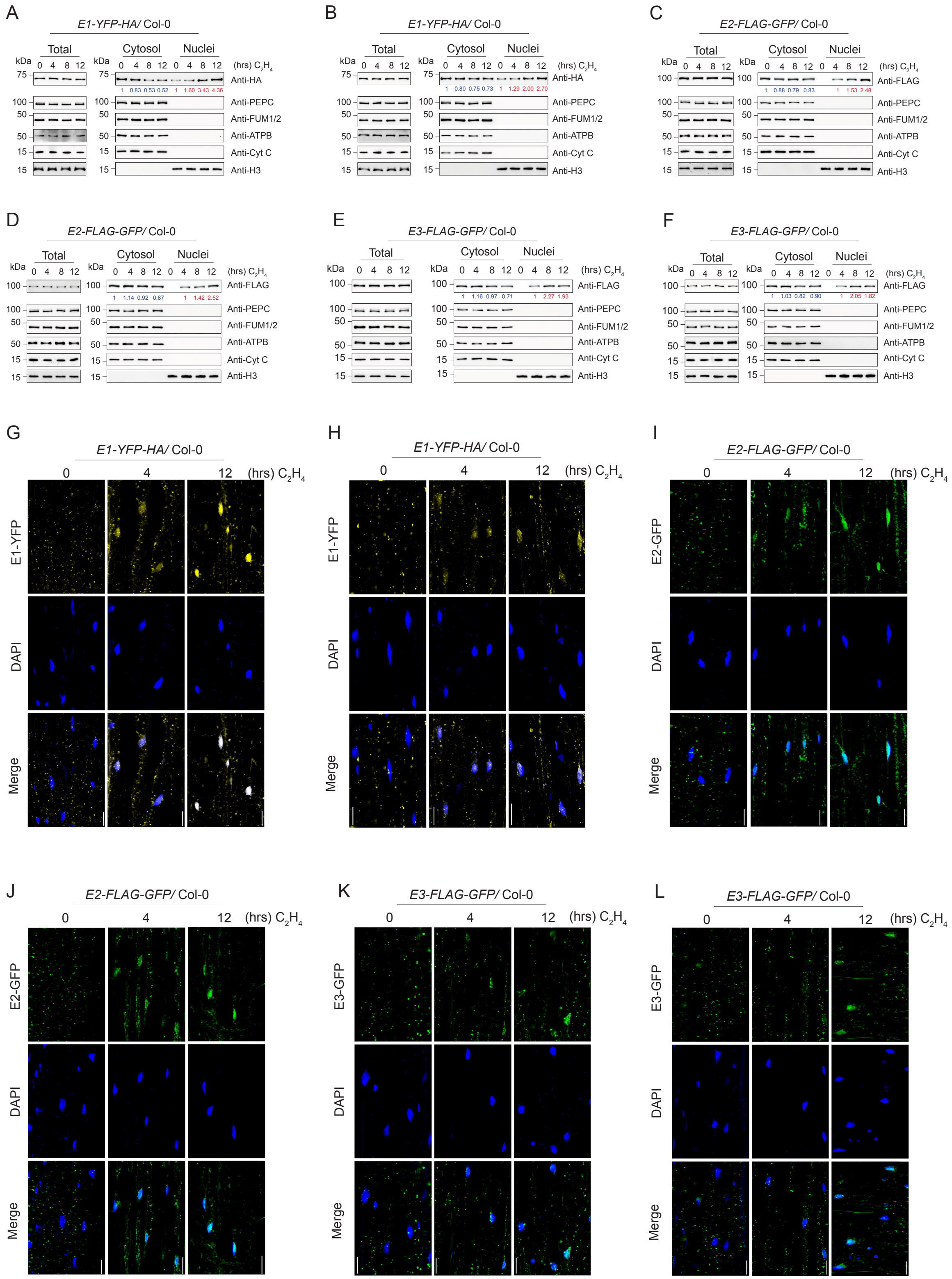
A
E1-YFP-HA/ Col-0
B
E1-YFP-HA/ Col-0
C
E2-FLAG-GFP/ Col-0
Total
Cytosol
Nuclei
(hrs) $C_2H_4$
kDa
Anti-HA
Anti-FLAG
Anti-PEPC
Anti-FUM1/2
Anti-ATPB
Anti-Cyt C
Anti-H3
D
E2-FLAG-GFP/ Col-0
E
E3-FLAG-GFP/ Col-0
F
E3-FLAG-GFP/ Col-0
G
E1-YFP-HA/ Col-0
H
E1-YFP-HA/ Col-0
I
E2-FLAG-GFP/ Col-0
E1-YFP
E2-GFP
E3-GFP
DAPI
Merge
J
E2-FLAG-GFP/ Col-0
K
E3-FLAG-GFP/ Col-0
L
E3-FLAG-GFP/ Col-0

**Supplemental Figure 2. Related to Figure 2. Nuclear PDC accumulation in response to ethylene.**

(**A**-**F**) Two additional independent biological replicates of fractionation immunoblots to examine the total and subcellular protein levels of PDC in of *E1-YFP-HA* (**A** and **B**), *E2-FLAG-GFP* (**C** and **D**), and *E3-FLAG-GFP* (**E** and **F**) transgenic plants with ethylene gas treatments of 0, 4h, 8h, and 12h. PDC E1 was probed with anti-HA antibody, and E2 and E3 were probed with anti-FLAG antibody in total protein extracts, cytoplasmic fractions, and nuclear fractions. PEPC, CytC, ATPB, FUM1/2, and histone H3 was used to evaluate the purities of nuclear and cytosolic fractionations, as well as to serve as loading controls. Blue number indicates PDC band intensity that normalized to PEPC signal. Red number indicates PDC band intensity that normalized to histone H3 signal. (**G**-**L**) Two additional independent biological replicate results of confocal microscopy images showing the subcellular localization of E1-YFP-HA (**G** and **H**), E2-FLAG-GFP (**I** and **J**), and E3-FLAG-GFP (**K** and **L**) with time series of ethylene gas treatments. DAPI staining labels nuclei. Scale bars is 20μm.

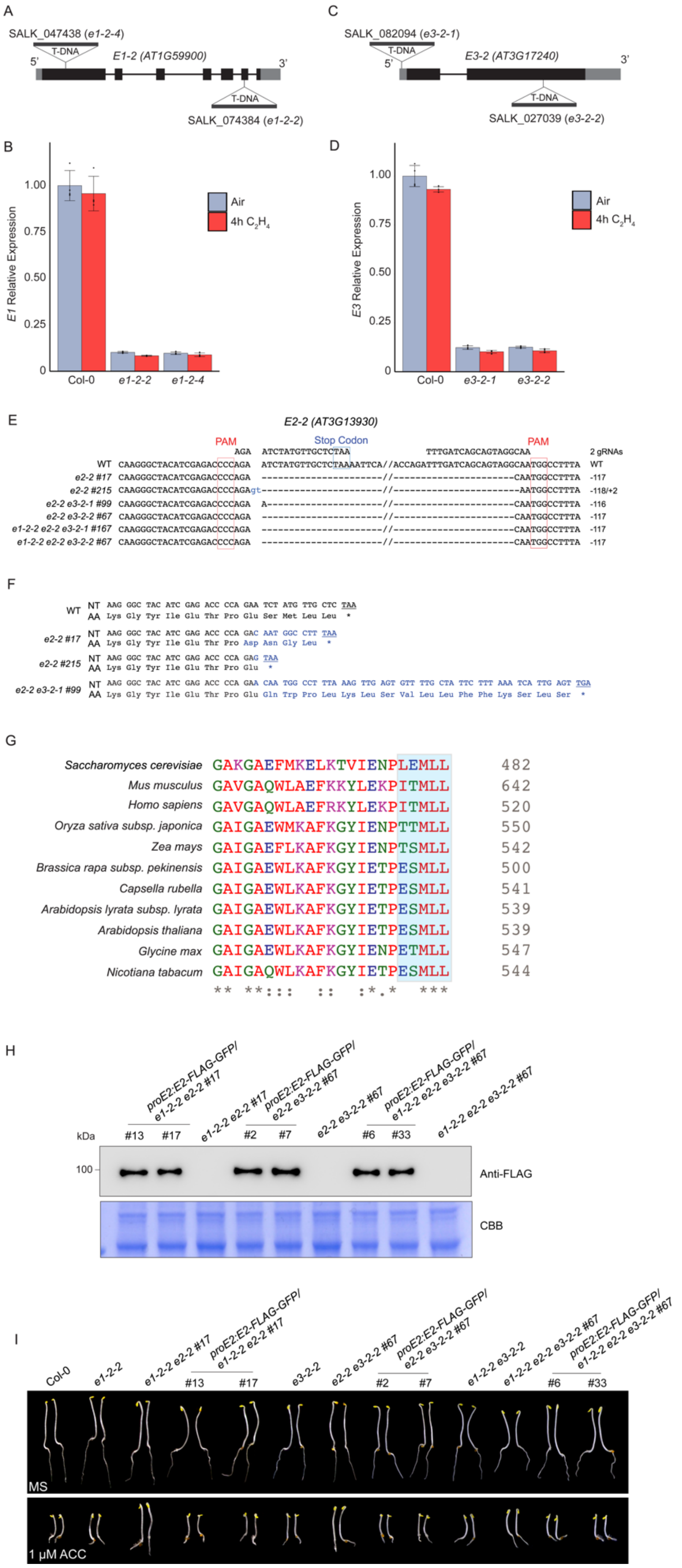
A
SALK_047438 (e1-2-4)
T-DNA
E1-2 (AT1G59900)
SALK_074384 (e1-2-2)
C
SALK_082094 (e3-2-1)
E3-2 (AT3G17240)
SALK_027039 (e3-2-2)
B
E1 Relative Expression
Air
4h C2H4
Col-0
e1-2-2
e1-2-4
D
E3 Relative Expression
e3-2-1
e3-2-2
E
E2-2 (AT3G13930)
PAM
Stop Codon
2 gRNAs
F
G
Saccharomyces cerevisiae
Mus musculus
Homo sapiens
Oryza sativa subsp. japonica
Zea mays
Brassica rapa subsp. pekinensis
Capsella rubella
Arabidopsis lyrata subsp. lyrata
Arabidopsis thaliana
Glycine max
Nicotiana tabacum
H
kDa
Anti-FLAG
CBB
I
MS
1 μM ACC

**Supplemental Figure 3. Related to Figure 3. Different *PDC* mutant alleles.**

(**A**) Diagram of the gene structure of *PDC E1* locus, and the T-DNA insertion loci of *e1-2-2* and *e1-2-4* were indicated. (**B**) qRT-PCR analysis of *E1* expression in 3-day-old etiolated seedlings of Col-0, *e1-2-2*, and *e1-2-4* with and without 4-hour ethylene treatments. Bar represents the average of relative expression of *E1* normalized to *Actin2* expression*.* Whisker represents ± SD (n =4). (**C**) Diagram to show the T-DNA insertion locus for *e3-2-1* and *e3-2-2* alleles. (**D**) qRT-PCR analysis of *E3* expression in the indicated 3-day-old etiolated seedlings with and without 4 hours of ethylene gas treatments. Values are average of relative expression of *E3* normalized to that of *Actin2;* error bar indicates the SD (n = 4). (**E**) DNA sequences to show the mutations in *e2* single mutants, *pdc* double or triple mutants that were generated by CRIPSR-Cas9. Deletions are shown as dashes. Red box indicates CRISPR gRNA PAM sequence. The numbers of deletion (base pair) for each mutation are shown on the right. (**F**) Potential mutated E2 protein amino acid sequences predicted from mutated *E2* nucleotide sequences. Mutated nucleotides and amino acids are in blue, and the stop codon is underlined and indicated by asterisk. The mutations of *E2* in *e2-2 #17* is the same as mutations of *E2* in *e2-2 e3-2-2 #67*, *e1-2-2 e2-2 e3-2-1 #167*, and *e1-2-2 e2-2 e3-2-2 #67*. (**G**) The alignment of protein sequence from E2 mutated region to show the evolutionary conservation across various species. Blue shade indicates the location of the varieties of mutations in *E2*. (**H**) Protein levels of *proE2:gE2-FLAG-GFP* in representative complemented lines from each genetic background. Coomassie blue staining was used as a loading control. (**I**) Representative 3-day-old etiolated seedlings grown on MS medium and 1 μM ACC containing MS medium showing the molecular complementation by *proE2:gE2-FLAG-GFP*.

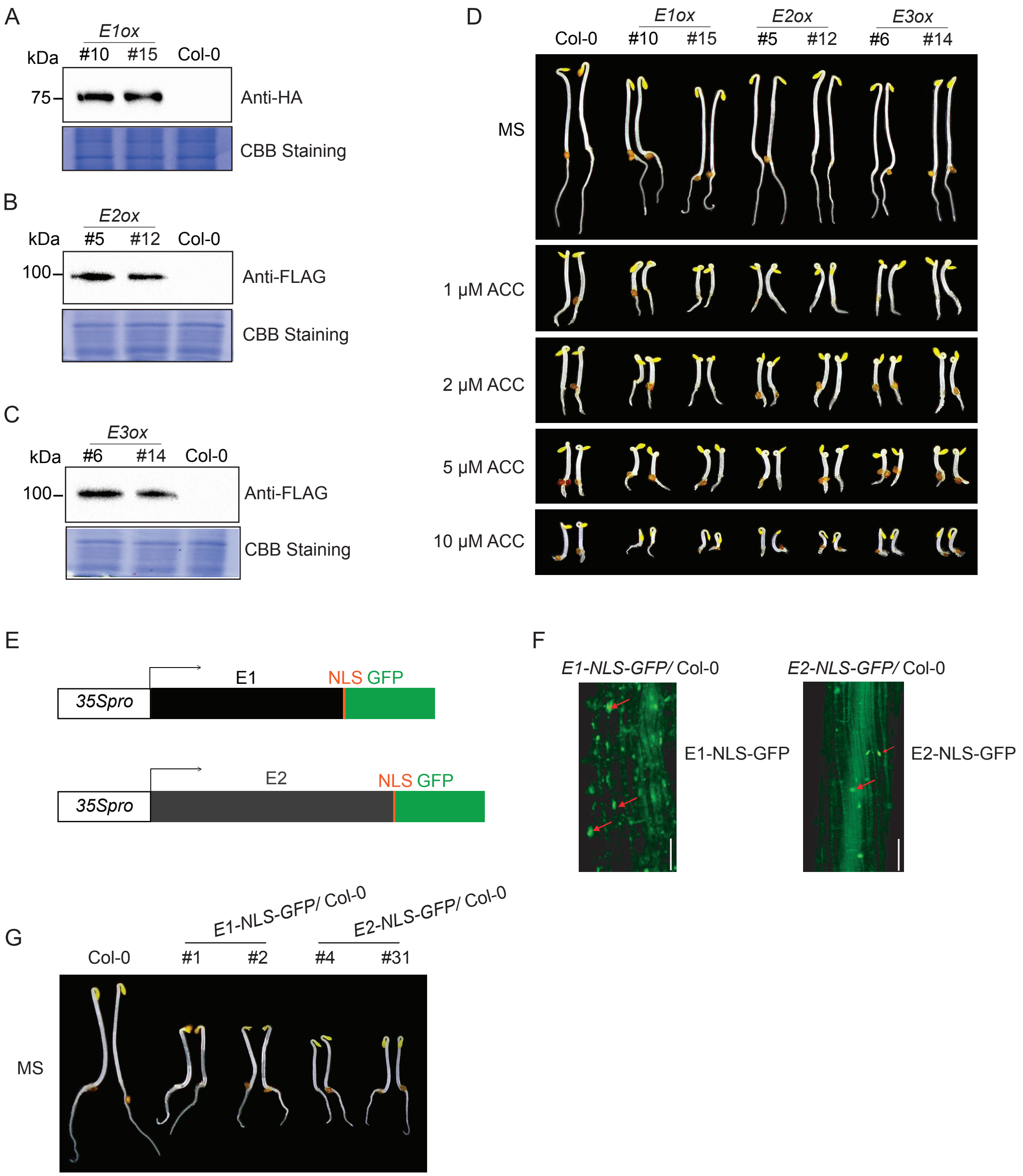

A
E1ox
kDa #10 #15 Col-0
75
Anti-HA
CBB Staining
B
E2ox
kDa #5 #12 Col-0
100
Anti-FLAG
CBB Staining
C
E3ox
kDa #6 #14 Col-0
100
Anti-FLAG
CBB Staining
D
Col-0
E1ox #10 #15
E2ox #5 #12
E3ox #6 #14
MS
1 μM ACC
2 μM ACC
5 μM ACC
10 μM ACC
E
35Spro
E1
NLS GFP
35Spro
E2
NLS GFP
F
E1-NLS-GFP/ Col-0
E1-NLS-GFP
E2-NLS-GFP/ Col-0
E2-NLS-GFP
G
Col-0
E1-NLS-GFP/ Col-0
#1 #2
E2-NLS-GFP/ Col-0
#4 #31
MS

**Supplemental Figure 4. Related to Figure 3. Analyses of *PDC* gain-of-function transgenic plants and *E1- NLS-GFP* and *E2-NLS-GFP* transgenic plants.**

(**A**-**C**) Western blot assays of E1, E2, and E3 protein levels in *E1ox* plants (**A**), *E2ox* plants (**B**), and *E3ox plants* (**C**) by anti-HA and anti-FLAG, respectively. Coomassie Brilliant Blue (CBB) Staining served as a loading control. (**D**) Photography of *E1ox*, *E2ox,* and *E3ox* seedings grown on MS medium containing 0, 1 μM, 2 μM, 5 μM, and 10 μM ACC in the dark for 3 days. (**E**) Diagrams of the binary vector containing chimeric *E1-NLS-eGFP* and *E2-NLS-eGFP* fused genes that are driven by the *35S* promoter. (**F**) Confocal image showing the nuclear localization of E1-NLS-eGFP and E2-NLS-eGFP in *E1-NLS-eGFP* and *E2-NLS-eGFP* transgenic plants without ethylene treatment. Red arrow indicates nuclei. Scale bar is 50μm. (**G**) Photography of *E1-NLS-GFP* and *E2-NLS-GFP* transgenic plants grown on MS medium without ACC in the dark for 3 days.

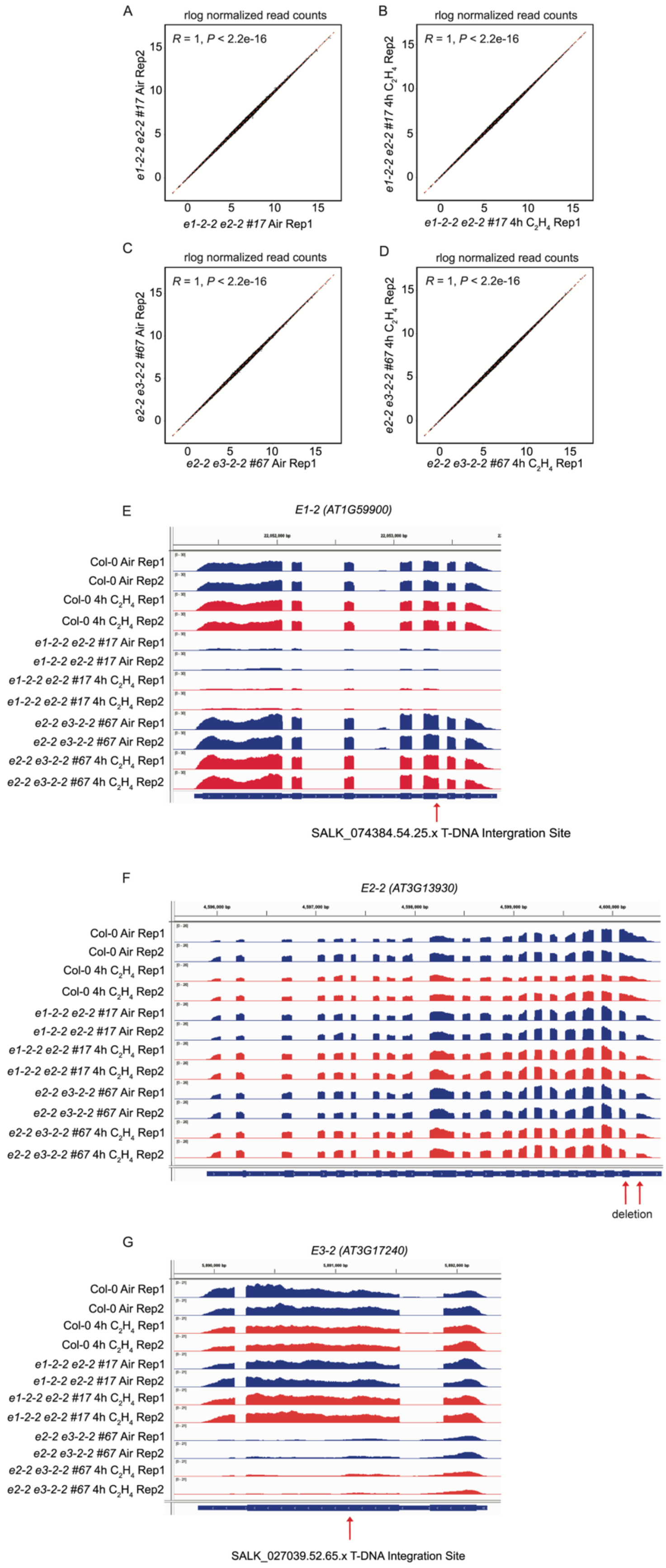
A
rlog normalized read counts
R = 1, P < 2.2e-16
e1-2-2 e2-2 #17 Air Rep2
e1-2-2 e2-2 #17 Air Rep1
B
rlog normalized read counts
R = 1, P < 2.2e-16
e1-2-2 e2-2 #17 4h C2H4 Rep2
e1-2-2 e2-2 #17 4h C2H4 Rep1
C
rlog normalized read counts
R = 1, P < 2.2e-16
e2-2 e3-2-2 #67 Air Rep2
e2-2 e3-2-2 #67 Air Rep1
D
rlog normalized read counts
R = 1, P < 2.2e-16
e2-2 e3-2-2 #67 4h C2H4 Rep2
e2-2 e3-2-2 #67 4h C2H4 Rep1
E
E1-2 (AT1G59900)
Col-0 Air Rep1
Col-0 Air Rep2
Col-0 4h C2H4 Rep1
Col-0 4h C2H4 Rep2
e1-2-2 e2-2 #17 Air Rep1
e1-2-2 e2-2 #17 Air Rep2
e1-2-2 e2-2 #17 4h C2H4 Rep1
e1-2-2 e2-2 #17 4h C2H4 Rep2
e2-2 e3-2-2 #67 Air Rep1
e2-2 e3-2-2 #67 Air Rep2
e2-2 e3-2-2 #67 4h C2H4 Rep1
e2-2 e3-2-2 #67 4h C2H4 Rep2
SALK_074384.54.25.x T-DNA Intergration Site
F
E2-2 (AT3G13930)
Col-0 Air Rep1
Col-0 Air Rep2
Col-0 4h C2H4 Rep1
Col-0 4h C2H4 Rep2
e1-2-2 e2-2 #17 Air Rep1
e1-2-2 e2-2 #17 Air Rep2
e1-2-2 e2-2 #17 4h C2H4 Rep1
e1-2-2 e2-2 #17 4h C2H4 Rep2
e2-2 e3-2-2 #67 Air Rep1
e2-2 e3-2-2 #67 Air Rep2
e2-2 e3-2-2 #67 4h C2H4 Rep1
e2-2 e3-2-2 #67 4h C2H4 Rep2
deletion
G
E3-2 (AT3G17240)
Col-0 Air Rep1
Col-0 Air Rep2
Col-0 4h C2H4 Rep1
Col-0 4h C2H4 Rep2
e1-2-2 e2-2 #17 Air Rep1
e1-2-2 e2-2 #17 Air Rep2
e1-2-2 e2-2 #17 4h C2H4 Rep1
e1-2-2 e2-2 #17 4h C2H4 Rep2
e2-2 e3-2-2 #67 Air Rep1
e2-2 e3-2-2 #67 Air Rep2
e2-2 e3-2-2 #67 4h C2H4 Rep1
e2-2 e3-2-2 #67 4h C2H4 Rep2
SALK_027039.52.65.x T-DNA Integration Site

**Supplemental Figure 5. Related to Figure 3. mRNA sequencing of PDC *e1-2-2 e2-2 #17* and *e2-2 e3-2-2 #67* mutants with air and ethylene treatments.**

(**A**-**D**) Scatter plots showing the reproducibility of *e1-2-2 e2-2 #17* and *e2-2 e3-2-2 #67* RNA-seq results with or without 4 hours of ethylene treatment. *r log* transformed raw read counts were plotted and used to calculate Pearson correlation coefficient (*R*). (**E**-**G**) IGV genome browser tracks of mRNA-seq to show *E1*, *E2*, and *E3* loci in the wild type and in the mutant plants before and after ethylene treatment. Red arrow indicates the T-DNA insertion site (**E** and **G**) or the CRISPR/Cas9 mediated deletion region (**F**).

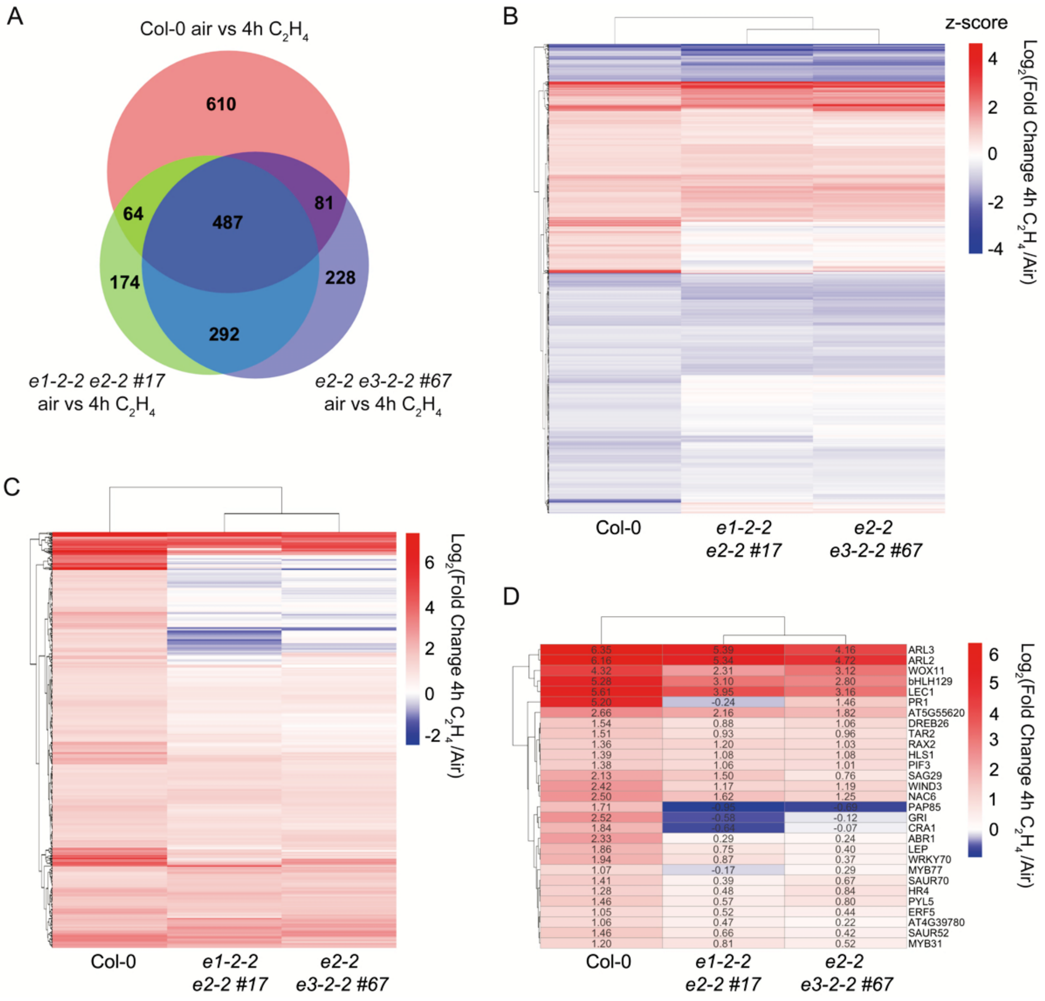


**Supplemental Figure 6. Related to Figure 3. PDC is required for ethylene response at transcriptional level.**

**(A)** Venn diagram to compare the ethylene regulated genes (|$\log_2$ FC| ≥ 1 and adjusted *P* value ≤ 0.05) in each double mutant and in Col-0. (**B**) Heatmap representation of z-score transformed $\log_2$(Fold Change $C_2H_4$/Air) of all ethylene-regulated differential expressed genes that were identified in Col-0 in *e1-2-2 e2-2 #17* and *e2-2 e3-2-2 #67* mutant backgrounds. (**C**) Heatmap to show the $\log_2$FC of genes that are up-regulated by ethylene in Col-0 but their up-regulation is compromised in the indicated plants. (**D**) Heatmap to compare the expression of part of typical ethylene up-regulated genes in Col-0 and in the indicated mutants. Raw $\log_2$FC value of each gene is annotated in the heatmap.

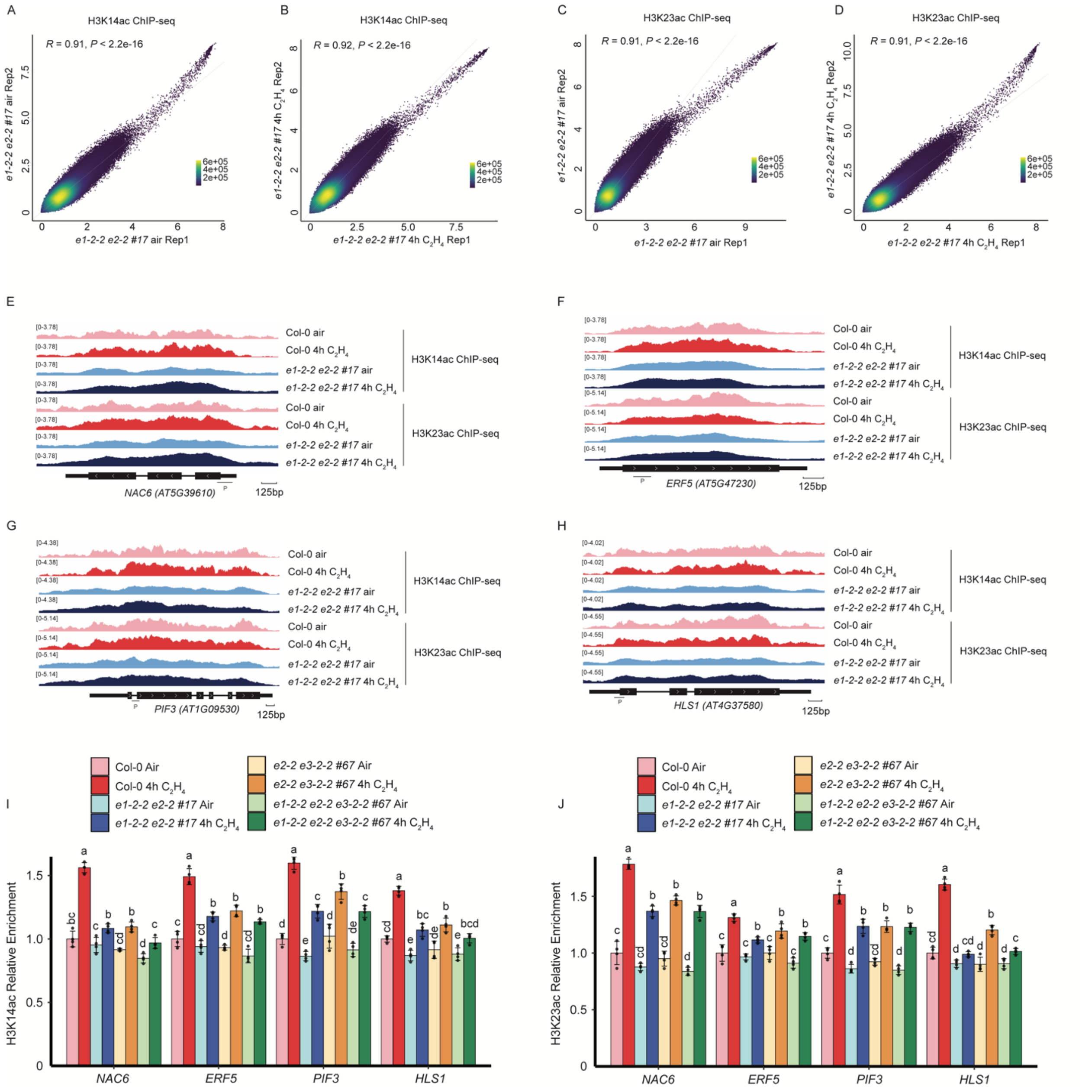


**Supplemental Figure 7. Related to Figure 4. ChIP sequencing of H3K14ac and H3K23ac in *pdc e1-2-2 e2-2 #17* mutants before and after ethylene treatment and ChIP-qPCR validation at selected ethylene responsive genes.**

**(A-D)** Scatter plots of mapped ChIP signals calculated by reads per genomic content (RPGC) showing the reproducibility of the replicates (Pearson correlation coefficient R > 0.9) from H3K14ac (**A** and **B**) and H3K23ac (**C** and **D**) ChIP sequencing. Each dot is a 100bp bin of the genome. Color key represents dot densities. (**E-H**) IGV genome browser snapshots to show

H3K14ac and H3K23ac ChIP signal from *NAC6* (**E**), *ERF5* (**F**), *PIF3* (**G**), and *HLS1* (**H**) loci. ChIP-seq signals were normalized by RPGC in Col-0 and *e1-2-2 e2-2 #17* under air and ethylene treatment. Horizontal lines represent the amplified regions for ChIP-qPCR and P represents ChIP-qPCR primer. Horizontal bracket represents scale bar of 125bp. (**I**-**J**) ChIP-qPCR to validate the enrichment of H3K14ac (**I**) and H3K23ac (**J**) in Col-0, *e1-2-2 e2-2 #17, e2-2 e3-2-2 #67,* and *e1-2-2 e2-2 e3-2-2 #67* etiolated seedlings treated with or without 4 hours of ethylene treatment of selected genes. Individual data point of the relative fold change normalized to Col-0 air is plotted as a dot. Different letters indicate significant differences ($P \leq 0.05$) between each genetic background and treatment condition calculated by a One-way ANOVA test followed by Tukey's HSD test.

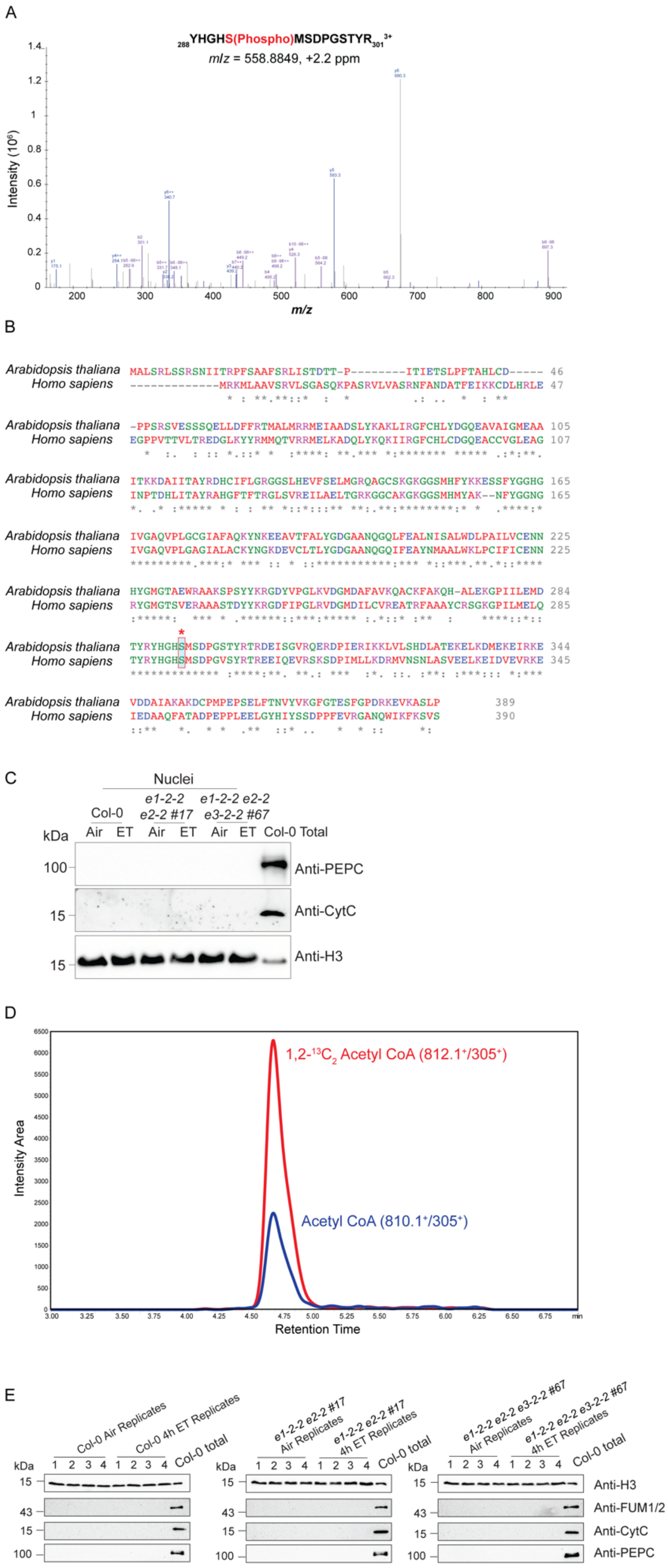
A
288YHGHS(Phospho)MSDPGSTYR301 3+
m/z = 558.8849, +2.2 ppm
Intensity (10^6)
m/z
B
Arabidopsis thaliana MALSRLSSRSNIITRPFSAAFSRLISTDTT-P---------ITIETSLPFTAHLCD----- 46
Homo sapiens -------------MRKMLAAVSRVLSGASQKPASRVLVASRNFANDATFEIKKCDLHRLE 47
Arabidopsis thaliana -PPSRSVESSSQELLDFFRTMALMRRMEIAADSLYKAKLIRGFCHLYDGQEAVAIGMEAA 105
Homo sapiens EGPPVTTVLTREDGLKYYRMMQTVRRMELKADQLYKQKIIRGFCHLCDGQEACCVGLEAG 107
Arabidopsis thaliana ITKKDAIITAYRDHCIFLGRGGSLHEVFSELMGRQAGCSKGKGGSMHFYKKESSFYGGHG 165
Homo sapiens INPTDHLITAYRAHGFTFTRGLSVREILAELTGRKGGCAKGKGGSMHMYAK--NFYGGNG 165
Arabidopsis thaliana IVGAQVPLGCGIAFAQKYNKEEAVTFALYGDGAANQGQLFEALNISALWDLPAILVCENN 225
Homo sapiens IVGAQVPLGAGIALACKYNGKDEVCLTLYGDGAANQGQIFEAYNMAALWKLPCIFICENN 225
Arabidopsis thaliana HYGMGTAEWRAAKSPSYYKRGDYVPGLKVDGMDAFAVKQACKFAKQH-ALEKGPIILEMD 284
Homo sapiens RYGMGTSVERAAASTDYYKRGDFIPGLRVDGMDILCVREATRFAAAYCRSGKGPILMELQ 285
Arabidopsis thaliana TYRYHGHSMSDPGSTYRTRDEISGVRQERDPIERIKKLVLSHDLATEKELKDMEKEIRKE 344
Homo sapiens TYRYHGHSMSDPGVSYRTREEIQEVRSKSDPIMLLKDRMVNSNLASVEELKEIDVEVRKE 345
Arabidopsis thaliana VDDAIAKAKDCPMPEPSELFTNVYVKGFGTESFGPDRKEVKASLP 389
Homo sapiens IEDAAQFATADPEPPLEELGYHIYSSDPPFEVRGANQWIKFKSVS 390
C
Nuclei
Col-0
e1-2-2 e2-2 #17
e1-2-2 e2-2 e3-2-2 #67
kDa
Air ET Air ET Air ET Col-0 Total
100
15
15
Anti-PEPC
Anti-CytC
Anti-H3
D
1,2-13C2 Acetyl CoA (812.1+/305+)
Acetyl CoA (810.1+/305+)
Intensity Area
Retention Time
min
E
Col-0 Air Replicates
Col-0 4h ET Replicates
Col-0 total
e1-2-2 e2-2 #17 Air Replicates
e1-2-2 e2-2 #17 4h ET Replicates
e1-2-2 e2-2 e3-2-2 #67 Air Replicates
e1-2-2 e2-2 e3-2-2 #67 4h ET Replicates
kDa
1 2 3 4 1 2 3 4
15
43
15
100
Anti-H3
Anti-FUM1/2
Anti-CytC
Anti-PEPC

**Supplemental Figure 8. Related to Figure 5. PDC E1 Ser292 phosphorylation site and 1,2-$^{13}C_2$ acetyl CoA isotope tracing experiment.**

(**A**) MS/MS spectrum of the peptide containing phosphorylated Ser292 of PDC E1. (**B**) Alignment of sequences of E1 from *Arabidopsis thaliana* and its homologue in *Homo sapiens*. Conserved Ser292 residue that its phosphorylation is inhibitory to E1 activity is circled and marked by a red asterisk. (**C**) Western blot analysis of PEPC, CytC, and histone H3 to evaluate the purify of nuclear purification and the input loading of samples used in pyruvate dehydrogenase activity assays in Fig. 5F. (**D**) LC-MS/MS detection of the difference between endogenous $^{12}C$ acetyl CoA (molecular weight: 810 g/mol) and 1,2-$^{13}C_2$ acetyl CoA (molecular weight: 812 g/mol) based on their molecular weights. Raw data from 3-day-old Col-0 etiolated seedlings treated with 4 hours of ethylene gas replicate 1 was shown. (**E**) Western blot analysis of PEPC, cytochrome C, FUM1/2, and histone H3 to evaluate the nuclear purification efficiencies for $^{13}C$ isotopic tracing experiments in Fig. 5H.

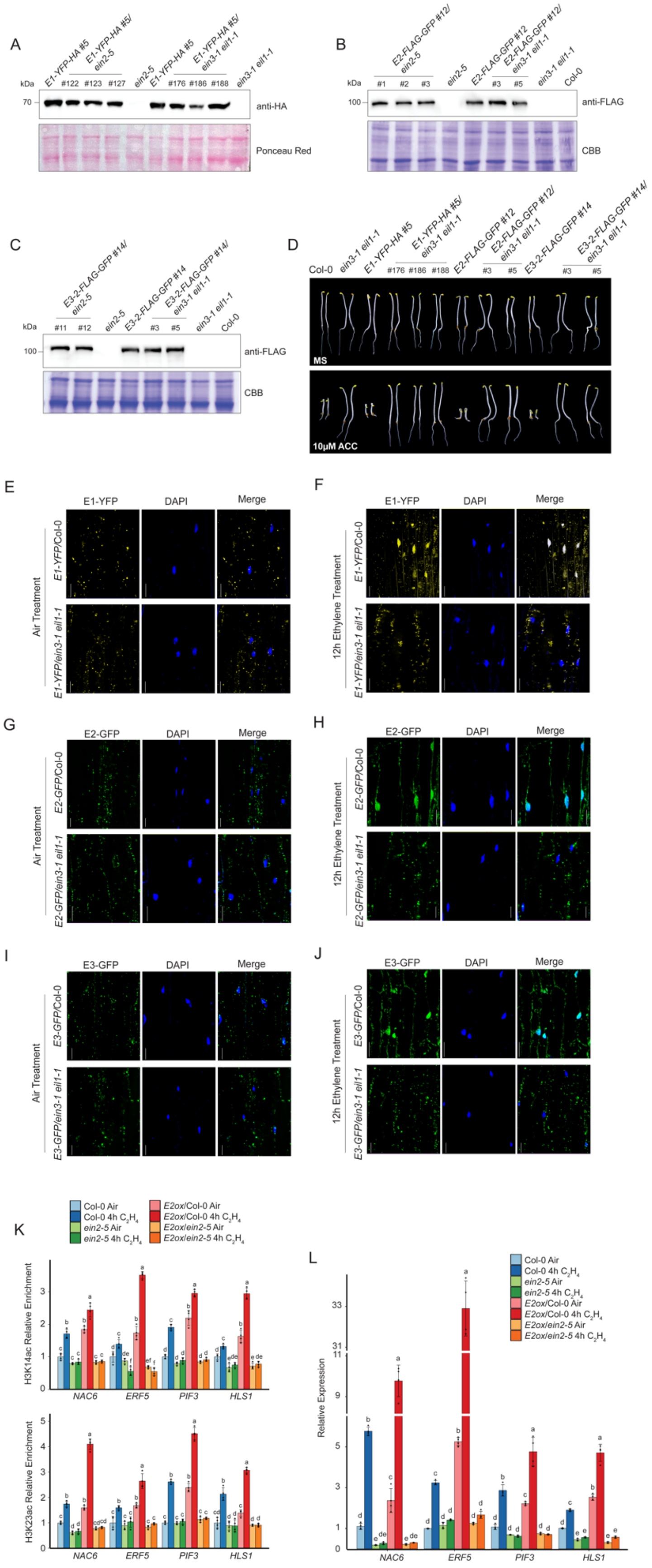

A
B
C
D
E
F
G
H
I
J
K
L
anti-HA
Ponceau Red
anti-FLAG
CBB
MS
10μM ACC
E1-YFP
DAPI
Merge
E2-GFP
E3-GFP
Air Treatment
12h Ethylene Treatment
H3K14ac Relative Enrichment
H3K23ac Relative Enrichment
Relative Expression
NAC6
ERF5
PIF3
HLS1

**Supplemental Figure 9. Related to Figure 6. Different assays to analyze the connection between PDC and *ein2-5* or *ein3-1eil1-1*.**

(**A**-**C**) Western blot assay of E1, E2 and E2 protein levels in *E1ox/ein2-5* and *E1ox/ein3-1 eil1-1* plants (**A**), *E2ox/ein2-5* and *E2ox/ ein3-1 eil1-1* plants (**B**), and *E3ox/ein2-5* and *E3ox/ ein3-1 eil1-1* plants (**C**) by anti-HA or anti-FLAG antibody. Ponceau Red staining or Coomassie Brilliant Blue (CBB) staining serves as a loading control. #122 of *E1ox/ein2-5*, #1 of *E2ox/ein2-5*, and #11 of *E3ox/ein2-5* were used for the following molecular assays. (**D**) Phenotypic analysis of PDC *E1ox*, *E2ox*, and *E3ox* in *ein3-1eil1-1* mutant. The seedlings were grown on MS medium supplemented with or without 10$\mu$M ACC in the continuous dark for 3 days before being photographed. (**E-J**) Subcellular localization of E1-YFP-HA (**E** and **F**), E2-FLAG-GFP (**G** and **H**), and E3-FLAG-GFP (**I** and **J**) in Col-0 or *ein3-1 eil1-1* mutant with air treatment or 12 hours of ethylene treatment, respectively. Scale bars, 20µm. (**K**) ChIP-qPCR analyses of selected target genes to evaluate H3K14ac (upper panel) and H3K23ac (lower panel) enrichment in Col-0, *E2ox, ein2-5,* and *E2ox/ein2-5* etiolated seedlings with or without 4 hours of ethylene treatment. Individual data point of the relative fold change to Col-0 air is plotted. Different letters indicate significant differences between each genotype and treatment condition calculated by a One-way ANOVA test followed by Tukey's HSD test with $P \leq 0.05$. (**L**) qRT-PCR analysis of target gene expressions in Col-0, *E2ox, ein2-5,* and *E2ox/ein2-5* 3-day-old etiolated seedlings with or without 4 hours of ethylene gas treatment. Values are average of relative expression from each gene normalized to that of *Actin2.* Error bar indicates the SD (n = 4) and different letters indicate statistically significant differences (One-way ANOVA test followed by Tukey's HSD test, $P \leq 0.05$).

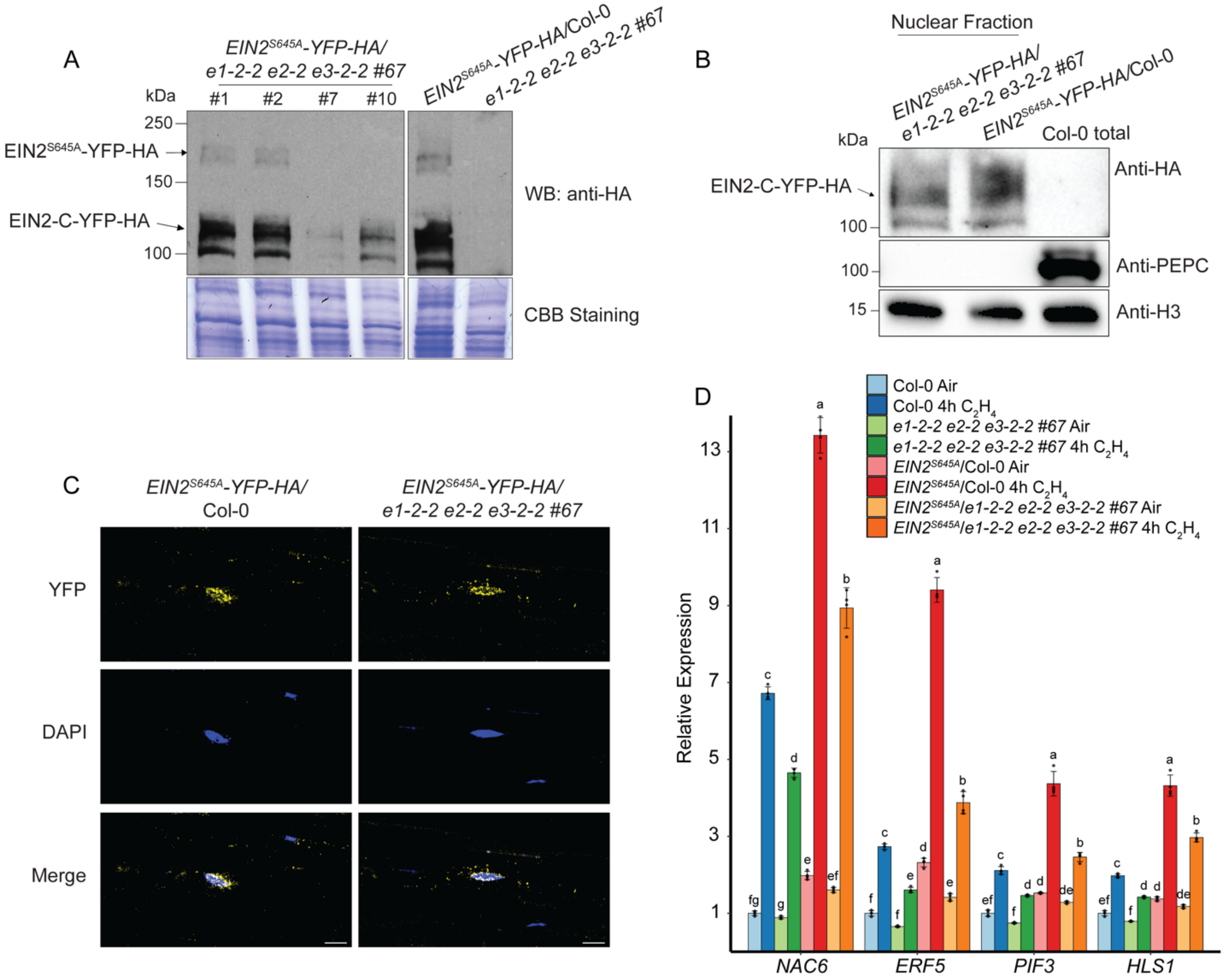


**Supplemental Figure 10. Related to Figure 6. EIN2$^{S645A}$ protein expression level, the nuclear accumulation of EIN2-C, and gene expression analysis of selected ethylene responsive genes in *EIN2$^{S645A}$/e1-2-2 e2-2 e3-3-2 #67*.**

(**A**) Western blot of EIN2$^{S645A}$ protein levels in *EIN2$^{S645A}$*/Col-0 and *EIN2$^{S645A}$/e1-2-2 e2-2 e3-3-2 #67* plants. Full-length of EIN2 and truncated EIN2-C were indicated by black arrows. Coomassie Brilliant Blue (CBB) staining was used as a loading control. #1 of *EIN2$^{S645A}$/e1-2-2 e2-2 e3-3-2 #67* was used for the following molecular experiments. (**B**) Western blot of nuclear EIN2-C protein levels in *EIN2$^{S645A}$/e1-2-2 e2-2 e3-3-2 #67* and *EIN2$^{S645A}$*/Col-0 after 4 hours of ethylene gas treatment. PEPC and histone H3 were used to mark cytosolic and nuclear proteins, respectively. (**C**) Confocal images to show the subcellular localizations of EIN2-C-YFP in Col-0 or *e1-2-2 e2-2* *e3-3-2 #67* treated with 4 hours of ethylene gas. Scale bars, 10μm. (**D**) qRT-PCR analysis to examine expression levels of target genes in 3-day-old etiolated seedlings of indicated genetic backgrounds treated with air or 4 hours of ethylene gas. Values are the average of relative expression from each target gene normalized to that of *Actin2*. Error bars represent the SD (n = 4). Different letters indicate statistically significant differences (One-way ANOVA test followed by Tukey's HSD test, $P \leq 0.05$).

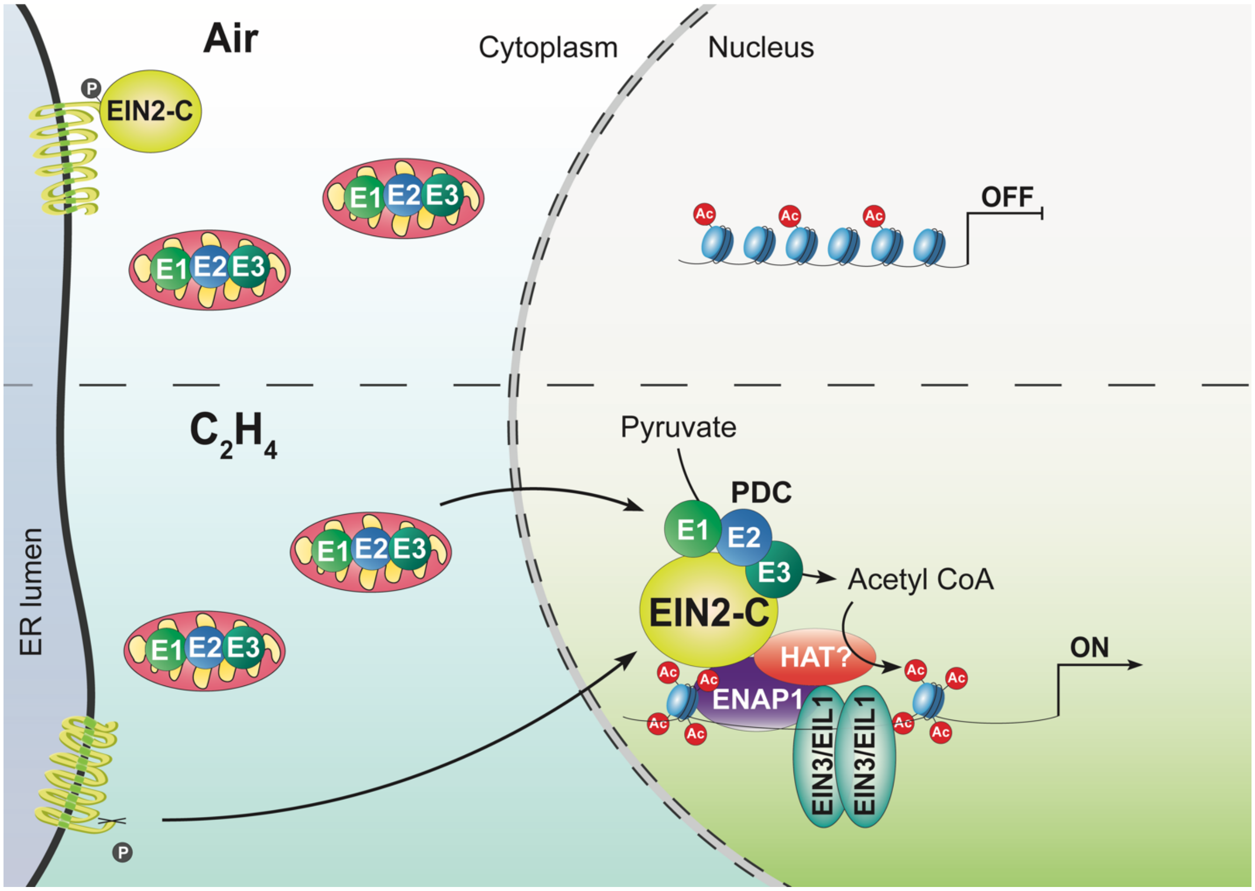


**Supplemental Figure 11. Proposed model of nuclear PDC in EIN2-directed histone acetylation in ethylene signaling.**

In the absence of ethylene, PDC resides in the mitochondria and EIN2 remains uncleaved on ER membrane (top). Upon ethylene exposure, cleaved EIN2-C is shuttled into the nucleus and PDC translocates from the mitochondria to the nucleus. By interacting with EIN2-C, nuclear PDC provides acetyl CoA to the EIN2-C-dependent histone acetylation machinery to elevate histone acetylation at H3K14 and H3K23 to regulate EIN3-dependent ethylene responsive transcriptional activation (bottom).

**Table S1. Primers used in the study**

| Primer Name | Sequence | Purpose |
|---|---|---|
| E1-Y2H-F | ACGCGTCGACAATGGCTCTATCACGCCTCT | Y2H construct |
| E1-Y2H-R | TCCCCCCGGGTCATGGAAGGGAAGCTTTGAC | Y2H construct |
| E2-Y2H-F | ACGCGTCGACAATGGCTTCTCGTATCATCAAT | Y2H construct |
| E2-Y2H-R | TCCCCCCGGGTTAGAGCAACATAGATTCTGGGG | Y2H construct |
| E3-Y2H-F | TCCCCCCGGGAATGGCGATGGCGAG | Y2H construct |
| E3-Y2H-R | CTAGACTAGTCTACATGTGAATGGGCTTGTC | Y2H construct |
| E1-pENTRY-pVP13-F | CACCTACCCATACGATGTTCCAGATTACGCTATGGCTCTATCACGCCTCT | *In vitro* expression |
| E1-pENTRY-pVP13-R | TCATGGAAGGGAAGCTTTGAC | *In vitro* expression |
| E2-pENTRY-pVP13-F | CACCGACTACAAAGACGATGACGACAAAATGGCTTCTCGTATCATCAAT | *In vitro* expression |
| E2-pENTRY-pVP13-R | TTAGAGCAACATAGATTCTGGGG | *In vitro* expression |
| E3-pMAL-pX2-F | CCGGAATTCGAACAAAAACTCATCTCAGAAGAGGATCTGATGGCGATGGCGAG | *In vitro* expression |
| E3-pMAL-pX2-R | ACGCGTCGACCTACATGTGAATGGGCTTGTC | *In vitro* expression |
| E1-pENTRY-pEarleyGate101-F | CACCATGGCTCTATCACGCCTCT | Binary vector |
| E1-pENTRY-pEarleyGate101-R | TGGAAGGGAAGCTTTGAC | Binary vector |
| E2-pCambia1300-F | CGGGGTACCATGGCTTCTCGTATCATCAAT | Binary vector |
| E2-pCambia1300-R | ACGCGTCGACGAGCAACATAGATTCTGGGG | Binary vector |
| E3-pCambia1300-F | CGGGGTACCATGGCGATGGCGAG | Binary vector |
| E3-pCambia1300-R | ACGCGTCGACCATGTGAATGGGCTTGTC | Binary vector |

| | | |
|---|---|---|
| nYFP-pEarleyGate101-F | GTGCCTAGGGTGAGCAAGGGCGAG | BiFC binary vector |
| nYFP-pEarleyGate101-R | CTAGACTAGTTTACTCGATGTTGTGGCGG | BiFC binary vector |
| cYFP-pCambia1300-F | ACGCGTCGACGACGGCAGCGTGC | BiFC binary vector |
| cYFP-pCambia1300-R | ACGCGAGCTCTTACTTGTACAGCTCGTCCATG | BiFC binary vector |
| NLS-BFP-F | ACGCGTCGACATGTTGAAGCGGTATAAACGTCGGTTAATGAGCGAAGAACTAATCAAGG | Binary vector |
| NLS-BFP-R | ACGCGAGCTCTTATAACCGACGTTTATACCGCTTCAATCCGCTCCCATTCAGC | Binary vector |
| LBb1.3 | ATTTTGCCGATTTCGGAAC | Genotyping |
| *e1-2-2*-GT-F | AATGATAGGGGAGCTTGTTGG | Genotyping |
| *e1-2-2*-GT-R | TGGTTCTTTCTGTGCAGTTGC | Genotyping |
| *e1-2-4*-GT-F | AGAACGCCGTAAGTAACCCTC | Genotyping |
| *e1-2-4*-GT-R | CTGAGAAAACCTCATGAAGCG | Genotyping |
| *e3-2-1*-GT-F | AGGATCGTGATCCAGCATATG | Genotyping |
| *e3-2-1*-GT-R | GAGATAACAACCCTCCCAAGG | Genotyping |
| *e3-2-2*-GT-F | CGGAAATTTTCTCCGATTTTC | Genotyping |
| *e3-2-2*-GT-R | CATCTTCTTCGGCTTTGTGAG | Genotyping |
| *e2-2*-CRISPR-DT1-BsF | ATATATGGTCTCGATTGTTAGAGCAACATAGATTCTGTT | E2 CRISPR-Cas9 |
| *e2-2*-CRISPR-DT1-F0 | TGTTAGAGCAACATAGATTCTGTTTTAGAGCTAGAAATAGC | E2 CRISPR-Cas9 |
| *e2-2*-CRISPR-DT1-R0 | AACTTGCCTACTGCTGATCAAACAATCTCTTAGTCGACTCTAC | E2 CRISPR-Cas9 |
| *e2-2*-CRISPR-DT1-BsR | ATTATTGGTCTCGAAACTTGCCTACTGCTGATCAAACAA | E2 CRISPR-Cas9 |
| *e2-2*-GT-F | AGTGGCTGAAAGCATTCAAG | Genotyping |
| *e2-2*-GT-R | TTTAAAGAATAGCAAAACACTCAACTT | Genotyping |
| Cas9-GT-F | CACCGACGAGTACAAGG | Genotyping |

| | | |
|---|---|---|
| Cas9-GT-R | GGCCCCTGAACTTAATCATG | Genotyping |
| E1-qRT-F | GTCACTCCATGTCTGATCCTG | Real time PCR |
| E1-qRT-R | TTTCTCGGTTGCTAGGTCATG | Real time PCR |
| E3-qRT-F | GTCTACACGTACCCTGAAGTTG | Real time PCR |
| E3-qRT-R | ATCTTGACCATTCCCTCTGC | Real time PCR |
| Actin2-qRT-F | CCCGCTATGTATGTCGC | Real time PCR |
| Actin2-qRT-R | AAGGTCAAGACGGAGGAT | Real time PCR |
| *proE2*-pCambia1300-F | AAAAGGCCTGCATGAGATAATGTTCTAATCTAAGACAT | Binary vector |
| *proE2*-pCambia1300-R | CGGGGTACCTGTTGTGCAATCGGAGC | Binary vector |
| E1-NLS-pCambia1300-F | CTAGTCTAGAATGGCTCTATCACGCCTCT | Binary vector |
| E1-NLS-pCambia1300-R | ACGCGTCGACTAACCGACGTTTATACCGCTTCAATGGAAGGGAAGCTTTGAC | Binary vector |
| E2-NLS-pCambia1300-F | CTAGTCTAGAATGGCTTCTCGTATCATCAAT | Binary vector |
| E2-NLS-pCambia1300-R | ACGCGTCGACTAACCGACGTTTATACCGCTTCAAGAGCAACATAGATTCTGGGG | Binary vector |
| PIF3-qRT-F | GCCATCGAGTATCTCAAGTCAC | Real time PCR |
| PIF3-qRT-R | AGGCATTCCCATACCCATTG | Real time PCR |
| NAC6-qRT-F | ACTCATAACTCACTACCTCAAACC | Real time PCR |
| NAC6-qRT-R | TTTTCTCCCATCTTAGCCTTCC | Real time PCR |
| HLS1-qRT-F | CGAATATCCACCCGAGTCATG | Real time PCR |
| HLS1-qRT-R | CGCTCCACGTACTTCTAACAG | Real time PCR |
| ERF5-qRT-F | TGTGACTGGGATTTAACGGG | Real time PCR |
| ERF5-qRT-R | CAACTGGGAATAACCAAACGG | Real time PCR |
| PIF3-ChIP-qPCR-F | CCGTGAGTCCCATTCACTTGTC | ChIP qPCR |
| PIF3-ChIP-qPCR-R | AGTTGATATCTGACCATTTTCCCA | ChIP qPCR |

| | | |
|---|---|---|
| NAC6-ChIP-qPCR-F | TAGAGAGGAGCTTCGTTGCTC | ChIP qPCR |
| NAC6-ChIP-qPCR-R | GTTTGGAGACGAAGAGGGAAG | ChIP qPCR |
| HLS1-ChIP-qPCR-F | CACCTTCCTCTCTATATATTAAACCCT | ChIP qPCR |
| HLS1-ChIP-qPCR-R | TCTAACCACCGTCATGTTTTGG | ChIP qPCR |
| ERF5-ChIP-qPCR-F | CTCCTAACGAAGTATCTGCACTTT | ChIP qPCR |
| ERF5-ChIP-qPCR-R | GATGATTCGTGCTTCATCCATG | ChIP qPCR |

**Table S2. Summary of total clean reads and aligned reads**

| **Sample Name** | **Repeat** | **Clean Reads** | **Aligned Reads** | **Align Rate (%)** |
|---|---|---|---|---|
| *e1-2-2 e2-2 #17* air (RNA-seq) | rep1 | 25600215 | 22932178 | 89.6 |
| *e1-2-2 e2-2 #17* air (RNA-seq) | rep2 | 28068692 | 25319409 | 90.2 |
| *e1-2-2 e2-2 #17* ethylene (RNA-seq) | rep1 | 31329229 | 28497496 | 91 |
| *e1-2-2 e2-2 #17* ethylene (RNA-seq) | rep2 | 30657907 | 27918368 | 91.1 |
| *e2-2 e3-2-2 #67* air (RNA-seq) | rep1 | 31892141 | 28828279 | 90.4 |
| *e2-2 e3-2-2 #67* air (RNA-seq) | rep2 | 40789675 | 37047373 | 90.8 |
| *e2-2 e3-2-2 #67* ethylene (RNA-seq) | rep1 | 30860626 | 28034354 | 90.8 |
| *e2-2 e3-2-2 #67* ethylene (RNA-seq) | rep2 | 31314332 | 28290889 | 90.3 |
| *e1-2-2 e2-2 #17* air input (ChIP-seq) | rep1 | 13888572 | 9746223 | 70.17 |
| *e1-2-2 e2-2 #17* air input (ChIP-seq) | rep2 | 16316376 | 11351075 | 69.57 |
| *e1-2-2 e2-2 #17* ethylene input (ChIP-seq) | rep1 | 15307947 | 10769023 | 70.35 |
| *e1-2-2 e2-2 #17* ethylene input (ChIP-seq) | rep2 | 17910315 | 12535083 | 69.99 |
| *e1-2-2 e2-2 #17* air H3K14ac (ChIP-seq) | rep1 | 23423131 | 17724647 | 75.67 |
| *e1-2-2 e2-2 #17* air H3K14ac (ChIP-seq) | rep2 | 19687063 | 14772192 | 75.04 |

| | | | | |
|---|---|---|---|---|
| *e1-2-2 e2-2 #17* ethylene H3K14ac (ChIP-seq) | rep1 | 19606552 | 14459614 | 73.75 |
| *e1-2-2 e2-2 #17* ethylene H3K14ac (ChIP-seq) | rep2 | 22993174 | 15838214 | 68.88 |
| *e1-2-2 e2-2 #17* air H3K23ac (ChIP-seq) | rep1 | 17236097 | 12295987 | 71.34 |
| *e1-2-2 e2-2 #17* air H3K23ac (ChIP-seq) | rep2 | 23249273 | 16405898 | 70.57 |
| *e1-2-2 e2-2 #17* ethylene H3K23ac (ChIP-seq) | rep1 | 24451561 | 17011539 | 69.57 |
| *e1-2-2 e2-2 #17* ethylene H3K23ac (ChIP-seq) | rep2 | 18938139 | 12157437 | 64.2 |
